# State-specific inhibition of NMDA receptors by memantine depends on intracellular calcium and provides insights into NMDAR channel blocker tolerability

**DOI:** 10.1101/2024.04.01.587624

**Authors:** Matthew B. Phillips, Nadya V. Povysheva, Karen A. Harnett-Scott, Elias Aizenman, Jon W Johnson

**Affiliations:** Department of Neuroscience and Center for Neuroscience, University of Pittsburgh, Pittsburgh, PA, USA; Department of Neurobiology and the Pittsburgh Institute for Neurodegenerative Diseases, University of Pittsburgh School of Medicine, Pittsburgh, PA, USA

## Abstract

NMDA receptors (NMDARs) are key mediators of neuronal Ca^2+^ influx. NMDAR-mediated Ca^2+^ influx plays a central role in synaptogenesis, synaptic plasticity, dendritic integration, and neuronal survival. However, excessive NMDAR-mediated Ca^2+^ influx initiates cellular signaling pathways that result in neuronal death and is broadly associated with neurological disease. Drugs targeting NMDARs are of great clinical interest, but widespread alteration of NMDAR activity can generate negative side effects. The NMDAR channel blocker memantine is a well-tolerated Alzheimer’s disease medication that shows promise in treatment of other neurological disorders. Memantine enhances desensitization of NMDARs in a subtype- and Ca^2+^-dependent manner, thereby more effectively inhibiting NMDARs on neurons that experience increased buildup of intracellular Ca^2+^. However, little is known about the properties or implications of the interaction between intracellular Ca^2+^ and NMDAR inhibition by memantine or other NMDAR channel blockers. Utilizing customized Ca^2+^ buffering solutions and whole-cell patch-clamp recordings, we demonstrated that memantine inhibition of both recombinant and native NMDARs increases with increasing intracellular Ca^2+^ and that the effect of intracellular Ca^2+^ on memantine action depends on NMDAR subtype. Neuroprotection assays and recordings of postsynaptic currents revealed that memantine preferentially inhibits NMDARs under neurotoxic conditions whereas ketamine, a clinically useful NMDAR channel blocker with strong side effects, inhibits strongly across contexts. Our results present a previously unexamined form of state-specific antagonism, Ca^2+^-dependent NMDAR channel block, that could have a profound impact on the design of drugs that selectively target NMDAR subpopulations involved in disease.

## Introduction

NMDA receptors (NMDARs) are ionotropic glutamate receptors that possess a unique set of biophysical properties, including dependence of gating on co-agonism, voltage-dependent block by Mg^2+^, high permeability to Ca^2+^, and slow gating kinetics (Mayer *et al*., 1984, 1987; Nowak *et al*., 1984; Johnson & Ascher, 1987; Vicini *et al*., 1998; Wyllie *et al*., 1998; Traynelis *et al*., 2010). This unique combination of properties allows NMDARs to control the magnitude and timing of Ca^2+^ influx during synaptic activity. Ca^2+^ influx due to NMDAR activity is vital to many aspects of neuronal function, including neuronal survival, synaptic development, and synaptic plasticity (Sheng *et al*., 1994; Malenka & Bear, 2004; Hardingham, 2006; Akgül & McBain, 2016).

The magnitude of NMDAR-mediated Ca^2+^ influx is a crucial determinant of the signaling pathways elicited by NMDAR activity. Low levels of NMDAR activity sustain small, prolonged increases of intracellular Ca^2+^ concentration ([Ca^2+^]_i_), which supports signaling cascades involved in synaptic depression. In contrast, intense, transient NMDAR activation leads to brief bouts of higher [Ca^2+^]_i_ and activates signaling cascades that drive synaptic potentiation (Lisman *et al*., 2002; Lüscher & Malenka, 2012). Sustained high NMDAR activity, however, initiates signaling cascades that result in neuronal death (Choi, 1987, 1992; Tymianski *et al*., 1993; Lau & Tymianski, 2010). Cell death elicited by excessive NMDAR-mediated Ca^2+^ influx, known as excitotoxicity, is a key feature of many nervous system disorders including Alzheimer’s disease, Alzheimer’s disease-related dementias, Huntington’s disease, and cell death following stroke, ischemia, and traumatic brain injury (Zorumski & Olney, 1993; Lipton, 1999, 2004; Hynd *et al*., 2004; Koutsilieri & Riederer, 2007; Dong *et al*., 2009; Olivares *et al*., 2012; Mota *et al*., 2014; Gardoni & Di Luca, 2015; Wang & Reddy, 2017, Gabrieli *et al*., 2021). Thus, NMDAR-mediated Ca^2+^ influx must be tightly regulated for proper nervous system function.

Although NMDARs are attractive targets for modulation by neurotherapeutic drugs, NMDAR pharmacology has proven remarkably complex. Due to the near-ubiquitous involvement of NMDARs in normal neuronal function, nonselective inhibition of NMDARs generates unacceptable side effects (Olney *et al*., 1989; Zorumski & Olney, 1993; Krystal *et al*., 1994; Muir, 2006). Most attempts to selectively target NMDAR subpopulations have focused on developing drugs that can distinguish between NMDAR subtypes. NMDARs display great subtype diversity, with subunits encoded by seven genes producing a single GluN1 subunit (with 8 distinct splice variants), four GluN2 subunits (A-D), and two GluN3 subunits (A and B) (Traynelis *et al*., 2010). The specific combination of subunits governs many NMDAR characteristics including subcellular localization, intracellular signaling partners, agonist affinity, gating kinetics, channel block, and pharmacology (Hardingham & Bading, 2010; Siegler Retchless *et al*., 2012; Paoletti *et al*., 2013; Hansen *et al*., 2014; Glasgow *et al*., 2015; Stroebel *et al*., 2018; Yi *et al*., 2018). However, subtype-selective antagonism has not yet produced clinically useful drugs, although some compounds have shown promise (Preskorn *et al*., 2008; Ibrahim *et al*., 2012).

The most clinically successful NMDAR antagonists are open channel blockers, drugs that bind in and prevent ion flux through open ion channels. However, most open channel blockers can elicit powerful and dangerous side effects, likely due to indiscriminate inhibition of NMDARs (Olney *et al*., 1989; Zorumski & Olney, 1993; Krystal *et al*., 1994). In stark contrast to other moderate and high-affinity NMDAR channel blockers, the adamantane derivative memantine is remarkably well-tolerated and can be administered chronically (Parsons, Danysz, & Quack, 1999; Chen & Lipton, 2006). Memantine is a clinically approved treatment for Alzheimer’s disease (Witt *et al*., 2004; Farlow *et al*., 2008; Mecocci *et al*., 2009; Danysz & Parsons, 2012) and shows promise in treating many other disorders including Parkinson’s disease, Alzheimer’s disease-related dementias, post-stroke cell death and dementia, schizophrenia, and disorders associated with rare *de novo* mutations of NMDAR subunits (Sonkusare *et al*., 2005; Lipton, 2006; Parsons *et al*., 2007; Berthier *et al*., 2009; Olivares *et al*., 2012; Pierson *et al*., 2014; Johnson *et al*., 2015; Di Iorio *et al*., 2017; Zheng *et al*., 2018; Folch *et al.,* 2018). A hypothesis regarding the clinical safety of memantine is that it may preferentially inhibit subpopulations of NMDARs involved in disease (Zhao *et al*., 2006; Léveillé *et al*., 2008; Okamoto *et al*., 2009; Xia *et al*., 2010), although the mechanism underlying this proposed selectivity has not yet been elucidated. Here, we present a mechanistic hypothesis to explain the clinical efficacy and safety of memantine: context-specific inhibition.

Memantine acts not only by blocking ion flux through NMDARs, but also by stabilizing a desensitized state of the NMDAR. Desensitized states are closed, agonist-bound states the occupancy of which depends both on NMDAR subtype and physiological context (Benveniste *et al*., 1990; Krupp *et al*., 1996, 1998; Villarroel *et al*., 1998; Glasgow *et al*., 2017; Phillips *et al*., 2020; Iacobucci and Pospescu 2024). We recently reported that memantine enhances NMDAR desensitization in a subtype-and context-specific manner (Glasgow *et al*., 2017). Memantine profoundly slows recovery of GluN1/2A receptors, but not GluN1/2B receptors, from desensitization. The effect of memantine on desensitization was absent in low extracellular Ca^2+^, and memantine inhibition of GluN1/2A receptors was shown to be more powerful in conditions supporting high levels of Ca^2+^ influx. These data suggest that memantine stabilizes a Ca^2+^-dependent desensitized state of the GluN1/2A receptor. Ca^2+^-dependent desensitization (CDD) acts as an endogenous negative-feedback loop by reducing NMDAR-mediated Ca^2+^ influx in response to increasing [Ca^2+^]_i_ (Legendre *et al*., 1993; Rozov & Burnashev, 2016; Iacobucci & Popescu, 2017, 2020). Stabilization of a Ca^2+^-dependent state by memantine offers a rational mechanism by which memantine can target specific NMDAR subpopulations involved in disease: preferential inhibition of NMDARs in neurons experiencing sustained high Ca^2+^ influx. Selective inhibition of overactive receptors is an ideal property of a well-tolerated drug, and elucidation of the relation between CDD and channel block of NMDARs may aid in the design of more efficacious pharmaceuticals.

Here we investigated the relation between NMDAR CDD and the mechanism of action of memantine. We found that memantine inhibition of recombinant and native GluN2A-containing NMDARs directly depends on [Ca^2+^]_i_. This [Ca^2+^]_i_ dependence of memantine inhibition requires occupancy of a specific Ca^2+^-dependent desensitized state only accessible by GluN2A-containing receptors. Using neuroprotection assays and recordings of evoked and postsynaptic currents to compare the effects of memantine and ketamine, a clinically useful NMDAR channel blocker with strong side effects, we found that memantine preferentially inhibits NMDARs under neurotoxic conditions while ketamine inhibits NMDARs uniformly across contexts, Our results present a previously uncharacterized form of state-specific NMDAR antagonism and strongly support the hypothesis that memantine acts as a context-specific antagonist.

## Methods

### Cell culture and transfection

Experiments were performed in tsA201 cell cultures (European Collection of Authenticated Cell Cultures) or primary cortical neuron cultures. tsA201 cells were maintained as previously described (Glasgow & Johnson, 2014) in Dulbecco’s modified Eagle’s medium (DMEM) supplemented with 10% fetal bovine serum and 1% GlutaMAX (Thermo Fisher Scientific). Cells were plated at a density of 10^5^ cells/dish in 35 mm petri dishes on 15 mm glass coverslips treated with poly D-lysine (0.1 mg/mL) and rat-tail collagen (0.1 mg/mL). 18-24 hours after plating, the cells were transfected using FuGENE 6 (Promega) with complementary DNA (cDNA) coding for enhanced green fluorescent protein (EGFP; Genbank ACS32473 in pCI-neo or pIRES) to identify transfected cells, WT rat GluN1-1a (GluN1; GenBank X63255 in pcDNA3.1 or U08261 in pCI-neo), and either GluN2A (GenBank M91561 in pcDNA1 or D13211 in pIRES), GluN2B (GenBank M91562 in pcDNA1), GluN2C (GenBank M91562 in pcDNA1), or GluN2D (GenBank L31611 in pcDNA1). EGFP was expressed using one of two plasmids: pCI-neo:EGFP:GluN1-1a or EGFP:pIRES:GluN2A, both kind gifts from Dr. Kasper Hansen. pCI-neo:EGFP:GluN1-1a was constructed by inserting cDNA encoding EGFP in pCI-neo under transcriptional control of the CMV promoter, between the CMV promoter and the GluN1 open reading frame (Yi *et al*., 2018). EGFP:pIRES:GluN2A was constructed by inserting cDNA encoding EGFP between the CMV promoter and the GluN2A open reading frame in pIRES, with the internal ribosome entry site (i.e., the IRES) between the EGFP and GluN2A open reading frames. Both plasmids allow co-expression of independent EGFP and NMDAR subunit proteins. For experiments with GluN1/2A receptors, cells were transfected with cDNA ratios of 1 GluN1: 1 GluN2. Cells were transfected with cDNA ratios of either 1 GluN1: 1 GluN2 or 1 GluN1: 2 GluN2 for experiments with GluN1/2B, GluN1/2C, and GluN1/2D receptors. 200 μM of the competitive NMDAR antagonist dl-APV was added to medium at the time of transfection to prevent NMDAR-mediated cell death.

Primary cortical cultures were prepared as previously described (Krall et al., 2020) from embryonic day 16 Sprague Dawley rats of both sexes. Pregnant rats (Charles River Laboratories) were sacrificed via CO_2_ inhalation. Brains of embryonic rats were dissected, and cortices were dissociated with trypsin. Dissociated neurons were then plated at a density of 6.6 * 10^5^ to 7.0 * 10^5^ cells/well on 15 mm glass coverslips in 6-well plates. Prior to plating, coverslips were acid-etched and treated with either poly-L-ornithine or poly-D-lysine. Non-neuronal cell proliferation was inhibited on DIV 15 by adding 1-2 μM cytosine arabinosine (AraC).

### Whole cell voltage-clamp recordings from cultured cells

Patch-clamp electrophysiological experiments were performed in the whole-cell voltage-clamp configuration. Recordings from tsA201 cells were performed 18-30 hours after transfection. Recordings from cultured neurons were performed after DIV 20 to allow for adequate GluN2A subunit expression, which occurs after roughly two weeks *in vitro* (Zhong *et al*., 1994; Li *et al*., 1998; Sinor *et al*., 2000). Pipettes were fabricated from borosilicate capillary tubing (outer diameter = 1.5 mm, inner diameter = 0.86 mm) using a Flaming Brown P-97 electrode puller (Sutter Instruments) and fire-polished to a resistance of 3.0 – 5.0 MΩ. For experiments in Figures 1 – 5 intracellular (pipette) solutions contained 120 – 130 mM CsCl, 10 mM HEPES, 4 mM MgATP and either 10 NTA, 10 HEDTA, or 10 BAPTA with the indicated concentrations of CaCl_2_ to give constant concentrations of free intracellular calcium (see *Intracellular solution preparation and determination of free [Ca^2+^] using the Ligand Optimization Method*). Internal solutions were pH balanced to 7.2 ± 0.05 with CsOH with a final osmolality of 290 ± 5 mOsM. For experiments measuring spontaneous NMDAR activity (Figure 6) a gluconate-based internal was used containing (in mM): 107 Cs-gluconate, 10 NaCl, 10 HEPES, 10 phosphocreatine, 4 MgATP, 0.3 GTP, and 10 BAPTA, balanced to pH 7.25 ± 0.05. Whole-cell currents were recorded with Axopatch 1D, Axopatch 200A, or MultiClamp 700A amplifiers and digitized using Digidata 1440A digitizers (Molecular Devices). Current signals were low-pass filtered at 5 kHz and sampled at 20 kHz using pCIamp10.3 or 10.7 (Molecular Devices). Series resistance was compensated between 85 – 90% in experiments for Figures 1-5, and data from cells with series resistance > 20 ΜΩ were excluded from analysis. Cells were clamped at −65 mV, accounting for empirically determined liquid junction potentials of −6 mV for Cl-based internals and −15 mV for gluconate-based internals.

**Figure 1.**
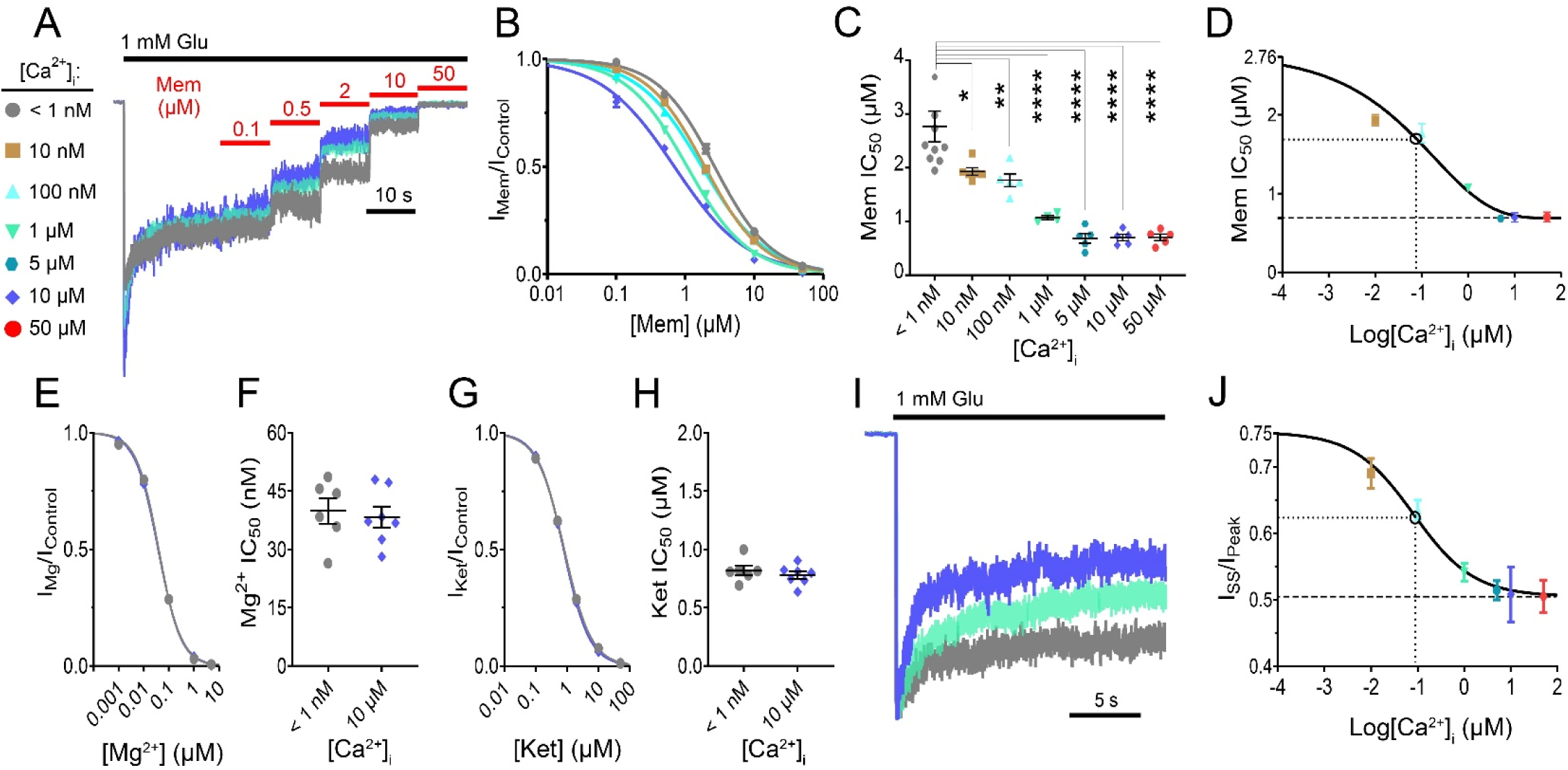
Ca^2+^-dependent block of GluN1/2A receptors by memantine. **A**, Overlay of WT GluN1/2A receptor currents used to measure memantine (Mem) concentration-inhibition curves during application of glutamate (Glu; black bar) and memantine (red bars) For visualization of differences in inhibition, only traces in conditions of [Ca^2+^]_i_ < 1 nM (gray), [Ca^2+^]_i_ = 1 μM (teal), and [Ca^2+^]_i_ = 10 μM (blue) are shown. [Ca^2+^]_e_ = 0.1 mM to prevent Ca^2+^ influx from altering [Ca^2+^]_i_. Currents are normalized to steady state current measured in 0 memantine. **B**, Memantine concentration-inhibition curves for [Ca^2+^]_i_ = 10 μM (blue), 1 μM (teal), 100 nM (light blue), 10 nM (gold), and < 1 nM (gray). Lines depict fit of Hill equation (**Equation 3**) to data. **C**, Summary of memantine IC_50_ values measured at the indicated [Ca^2+^]_i_ (< 1 nM Ca^2+^ : IC_50_ = 2.76 ± 0.27 μM, n = 9; 10 nM Ca^2+^ : IC_50_ = 1.93 ± 0.07 μM, n = 6; 100 nM Ca^2+^ : IC_50_ = 1.76 ± 0.12 μM, n = 5; 1 μM Ca^2+^ : IC_50_ = 1.07 ± 0.04 μM, n = 4; 5 μM Ca^2+^ : IC_50_ = 0.69 ± 0.06 μM, n = 5; 10 μM Ca^2+^ : IC_50_ = 0.69 ± 0.05 μM, n = 5; 50 μM Ca^2+^ : IC_50_ = 0.70 ± 0.06 μM, n = 5). ANOVA with Sidak’s post hoc test. *p < 0.05, **p < 0.01, ****p < 0.0001. **D,** Curve describing the effect of [Ca^2+^]_i_ on memantine IC_50_. Line depicts fit of **Equation 4** to data. Memantine becomes more potent (IC_50_ decreases) as [Ca^2+^]_i_ increases. Dashed line at memantine IC_50_ = 0.69 μM depicts minimum IC_50_. Dotted lines and open circle show the [Ca^2+^]_i_ required to induce a half-maximal effect on memantine IC_50_ ([Ca^2+^]_i_ = 54 nM). **E**, Mg^2+^ concentration-inhibition curves for [Ca^2+^]_i_ < 1 nM (gray) and [Ca^2+^]_i_ = 10 μM (blue). Lines depict fit of **Equation 3** to data. **F**, Summary of Mg^2+^ IC_50_ values measured at [Ca^2+^]_i_ of < 1 nM (gray; IC_50_ = 39.9 ± 3.3 nM, n = 6) and 10 μM (blue; IC_50_ = 38.3 ± 2.7 nM, n = 7). 2-tailed Student t-test, p = 0.72. **G**, Ketamine concentration-inhibition curves for [Ca^2+^]_i_ < 1 nM (gray) and [Ca^2+^]_i_ = 10 μM (blue). Lines depict fit of **Equation 3** to data. **H**, Summary of ketamine IC_50_ values measured at [Ca^2+^]_i_ of < 1 nM (gray; IC_50_ = 0.82 ± 0.04, n = 6) and 10 μM (blue; IC_50_ = 0.78 ± 0.03 μM, n = 7). 2-tailed Student t-test, p = 0.46. **I**, Overlay of WT GluN1/2A receptor currents used to measure effect of [Ca^2+^]_i_ on desensitization; recordings shown are same as A, with currents instead normalized to peak response to Glu. **J**, Curve describing the effect of [Ca^2+^]_i_ on desensitization. Line depicts fit of **Equation 4** to data, dashed line at I_SS_/I_Peak_ = 0.505 depicts maximum desensitization, dotted lines and open circle show [Ca^2+^]_i_ required to induce a half-maximal effect on desensitization ([Ca^2+^]_i_ = 82 nM). For **C**, **F**, and **H**, points represent values from individual cells, bars and error bars depict mean ± SEM. For **B**, **D**, **E**, **G**, and **J**, data are depicted as mean ± SEM and some error bars are smaller than symbols. [Ca^2+^]_T_ and buffer used for each internal solution are given in **Table 2**, and [B]_T_ for each buffer is given in **Table 1**.

**Table 1.** Ligand Optimization Method parameters.

| Optimized Parameter | BAPTA | HEDTA | NTA |
| --- | --- | --- | --- |
| s (mV/decade $\text{Ca}^{2+}$) | 28.7 | 28.8 | 28.3 |
| $\Sigma$ | $8.47 * 10^{-9}$ | $7.64 * 10^{-18}$ | $1.79 * 10^{-16}$ |
| $[\text{B}]_{\text{T}}$ (M) | $9.49 * 10^{-3}$ | $9.04 * 10^{-3}$ | $8.71 * 10^{-3}$ |
| $K_{\text{d}}$ (M) | $1.44 * 10^{-7}$ | $2.24 * 10^{-6}$ | $8.15 * 10^{-5}$ |
| $E^0$ (mV) | 57.4 | 57.5 | 56.5 |

**Table 2.** Measured [Ca^2+^]F values in Ca^2+^ Buffer solutions.

| Target $[\text{Ca}^{2+}]_{\text{F}}$<br>(M) | Buffer | $[\text{Ca}^{2+}]_{\text{T}}$ (M) | Predicted $\Delta E$<br>(mV) | Measured $\Delta E$<br>(mV) | Measured $[\text{Ca}^{2+}]_{\text{F}}$<br>(M) |
| --- | --- | --- | --- | --- | --- |
| $1 * 10^{-8}$ | BAPTA | $6.16 * 10^{-4}$ | -164.58 | -164.6 | $9.98 * 10^{-9}$ |
| $1 * 10^{-7}$ | BAPTA | $3.89 * 10^{-3}$ | -142.51 | -142.5 | $1.00 * 10^{-7}$ |
| $1 * 10^{-6}$ | HEDTA | $2.79 * 10^{-3}$ | -115.07 | -115.1 | $9.98 * 10^{-7}$ |
| $5 * 10^{-6}$ | HEDTA | $6.20 * 10^{-3}$ | NA | -95.3 | $4.87 * 10^{-6}$ |
| $1 * 10^{-5}$ | HEDTA | $7.39 * 10^{-3}$ | -86.30 | -86.3 | $1.00 * 10^{-5}$ |
| $5 * 10^{-5}$ | NTA | $3.36 * 10^{-3}$ | -64.99 | -65.0 | $5.01 * 10^{-5}$ |

#5 $\mu\text{M}$ solution prepared using MAXCHELATOR predictions. All other solutions prepared using LOM predictions. All $[\text{Ca}^{2+}]_{\text{F}}$ values measured using LOM after final solution preparation.

Control bath solution (referred to as external solution) for tsA201 cell experiments contained (all concentrations in mM): 140 NaCl, 2.8 KCl, 10 HEPES, 0.01 EDTA, 0.1 glycine, and either 0.1 or 1 CaCl_2_. For cultured neuron experiments, external solution contained either 140 NaCl, 2.8 KCl, 10 HEPES, 0.01 EDTA, 0.01 glycine (glycine lowered from tsA201 cell experiments to prevent activation of inhibitory glycine receptors), and 0.1 CaCl_2_ (for Figure 5A-D experiments), or 126 NaCl, 2.5 KCl, 1.25 NaH_2_PO_4_, 1 CaCl_2_, 24 NaHCO_3_, 10-20 glucose, and 0.01 glycine (for Figure 6C-K experiments). 0.2 - 1 μM tetrodotoxin (TTX) was used to prevent synaptic events or action potential escape. Agonists and antagonists were added to external solutions on the day of experiments. 1 mM glutamate (diluted from 1 M stock) was used for tsA201 cell experiments, and 10 μM NMDA (diluted from 10 mM stock) for neuronal experiments. The indicated concentrations of memantine and ketamine (both diluted from 10 mM stocks in dH_2_O) were used for both tsA201 cell and neuronal experiments. Control, agonist, and antagonist solutions were delivered to the patched cell via ten polyimide barrels using our in-house fabricated rapid-switching fast perfusion system (Blanpied *et al*., 1997; Glasgow & Johnson, 2014). Solutions changes were performed by moving the barrel position relative to the patched cell with a voice-coil motor controlled by a custom program. Solution flow rate was maintained at 1 – 2 mL/min for all experiments utilizing the fast perfusion system. NMDAR-mediated miniature postsynaptic currents (mEPSCs) were isolated using gabazine (10 µM; Ascent Scientific), 2,3-dihydroxy-6-nitro-7-sulfamoylbenzo(F)quinoxaline (NBQX; 20 µM; Ascent Scientific). For all experiments manipulating internal calcium concentration except those shown in Figure 4, data was collected 5 – 10 min after break-in.

### Whole cell voltage-clamp recordings from prefrontal cortex brain slices

Electrophysiological experiments were performed in brain slices from 2- to 3-month-old wild-type C57BL/6J male mice. All animal procedures were in accord with the National Institutes of Health’s Guide for the Care and Use of Laboratory Animals and approved by the University of Pittsburgh Institutional Animal Care and Use Committee. Brain slices were prepared as previously described (Glasgow et al., 2017). Briefly, mice were thoroughly anesthetized with chloral hydrate. After decapitation, the brain was quickly removed and placed in ice-cold artificial CSF (ACSF) bubbled with a 95% O_2_/5% CO_2_ gas mixture. The front half of the brain containing the prelimbic cortex was excised for slicing. ACSF used for slicing and incubation contained (in mM): 126 NaCl, 2.5 KCl, 1.25 NaH_2_PO_4_, 1 MgSO_4_, 2 CaCl2, 24 NaHCO_3_, and 10-20 glucose (pH 7.25-7.3). Coronal slices (350 µm thick) were cut with a vibratome (VT1000S; Leica). Slices were incubated at 37°C for ∼1 h and further stored at room temperature until they were transferred to a recording chamber containing circulating ACSF bubbled with a 95% O_2_/5% CO_2_ gas mixture at 31-32°C. Electrodes for whole-cell recordings were pulled as described earlier to a resistance of 5-10 ΜΩ. Pipettes were filled with either low [Ca^2+^] or high [Ca^2+]^ internal solutions. Low-[Ca^2+^] (< 1nM Ca^2+^) internal contained (in mM): 107 Cs-gluconate, 10 NaCl, 10 HEPES, 10 phosphocreatine, 4 MgATP, 0.3 GTP, and 10 BAPTA. High-[Ca^2+^] (∼50 µM Ca^2+^) internal contained (in mM): 100 Cs-gluconate, 10 NaCl, 10 HEPES, 10 phosphocreatine, 4 MgATP, 0.3 GTP, 3.4 CaCl_2_, and 10 NTA. Both internals were balanced to pH 7.25 ± 0.05 with CsOH. Recordings were performed in ACSF containing: 126 NaCl, 2.5 KCl, 1.25 NaH_2_PO_4_, 1 CaCl_2_, 0.5 MgSO_4_, 24 NaHCO_3_, 10-20 glucose, and 0.01 glycine (pH 7.25-7.3). NMDAR-mediated postsynaptic currents (NMDAR-EPSCs) were isolated with gabazine (10 µM; Ascent Scientific) and 2,3-dihydroxy-6-nitro-7-sulfamoylbenzo(F)quinoxaline (NBQX; 20 µM; Ascent Scientific).

Pyramidal neurons in prefrontal cortical layers II/III were visualized by IR-DIC optics using an Axioskop microscope (Carl Zeiss), digital camera (CoolSnap; Photometrics), and 60X-water-immersion objective. The neurons were identified based on their characteristic triangular soma and apical dendrites. Whole-cell postsynaptic currents were recorded using a Multi-Clamp 700A amplifier (Molecular Devices), low-pass filtered at 2 kHz, and sampled at 10 kHz using a Digidata 1440 digitizer and Clampex 10.2 software (Molecular Devices). Series resistance compensation was not used. Access resistance was measured to be 10-20 MΩ and remained stable during experiments (±30% change) for the analyzed neurons. Neurons were clamped at −65 mV, correcting for an empirically determined junction potential of −15 mV. NMDAR*-*EPSCs were evoked by extracellular stimulation with theta-glass bipolar electrodes placed on the border of white matter and layer VI. Stimulation currents were generated with A360 stimulus isolator (World Precision Instruments) and were triggered digitally with Clampex. NMDAR-EPSCs were evoked by applying trains of 5 stimuli at 25 Hz (40 ms interstimulus intervals) with an intertrain interval of 10 s.

### Intracellular solution preparation and determination of free [Ca^2+^] using the Ligand Optimization Method

All intracellular (pipette) solutions contained 120 – 130 mM CsCl, 10 mM HEPES, and 4 mM MgATP and were pH balanced to 7.2 ± 0.05 with CsOH. To allow for study of the effects of known, constant [Ca^2+^]_i_ on channel block, each intracellular solution also contained empirically determined concentrations of CaCl_2_ and BAPTA, HEDTA, or NTA to buffer Ca^2+^ to the desired concentrations. Because estimation of free Ca^2+^ concentrations ([Ca^2+^]_F_) in buffered solutions are subject to multiple sources of error (McGuigan *et al*., 2016), we utilized the Ligand Optimization Method (LOM (McGuigan *et al*., 1991, 2006)) to (1) aid in the design of intracellular solutions containing known concentrations of buffered [Ca^2+^]_F_ and (2) empirically determine [Ca^2+^]_F_ following solution preparation. The LOM is a multi-step process that obtains the best fit of the Nicolsky-Eisenman equation (Nicolsky *et al*., 1967) to data measured with a Ca^2+^-selective electrode by optimizing four parameters vital to accurate determination of [Ca^2+^]_F_: the slope of the electrode at [Ca^2+^]s < 10 μM (*s*), the lumped interference constant (Σ) describing the nonlinearity of the electrode at low [Ca^2+^]_F_, the total concentration of the Ca^2+^ binding buffer ([B]_T_), and the K_d_ (equilibrium dissociation constant for binding of Ca^2+^ and buffer).

All solutions used for the LOM were prepared from a background solution containing 120 mM CsCl and 10 mM HEPES and balanced to pH 7.2 with CsOH. Ionic content of the background solution was designed to mimic our typical intracellular solutions. Seven calibration solutions, necessary for determination of *s*, were prepared by adding CaCl_2_ to background solution (without Ca2+ buffer) to produce total [Ca^2+^] ([Ca^2+^]_T_) ranging from 0.5 - 10 mM. 10 Ca^2+^ buffer solutions containing 10 mM of the calcium chelators BAPTA, HEDTA, or NTA (measured by weight) and known concentrations of [Ca^2+^]_T_ were prepared from background solution using the ratiometric method (McGuigan *et al*., 2014). All measurements of [Ca^2+^]_F_ were made at 25° C using a Ca^2+^-selective combination electrode (Orion 9720BNWP, ThermoFisher) and a pH meter in mV mode (Accument AR15, ThermoFisher). To obtain values for fitting of the Nicolsky-Eisenman equation, we first measured electrical potentials of the calibration solutions in order of descending [Ca^2+^]_T_ and then measured of electrical potentials of the Ca^2+^ buffer solutions in order of descending [Ca^2+^]_T_. Relative potentials (ΔE) for each solution were then calculated for each solution by subtracting the potential measured in the [Ca^2+^]_T_ = 10 mM calibration solution. The relative potentials were then fit using CaSALE, a custom R program written specifically for use with the LOM (kindly provided by Dr. James Kay; (McGuigan *et al*., 2014)) that automates the iterative optimization of *s*, Σ, [B]_T_, and K_d_. Values of optimized parameters are listed in Table 1.

After LOM curves were obtained for each Ca^2+^ buffer, [Ca^2+^]_T_s required to give desired [Ca^2+^]_F_ in our intracellular solutions were calculated using the optimized [B]_T_, and K_d_ values with **Equation 1**:

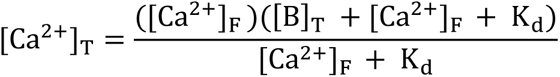

Buffers with K_d_ closest to the target [Ca^2+^]_F_ were used for each solution: BAPTA (K_d_ = 144 nM) for [Ca^2+^]_F_ < 1 μM, HEDTA (K_d_ = 2.24 μM) for [Ca^2+^]_F_ 1 – 10 μM, and NTA (K_d_ = 81.5 μM) for [Ca^2+^]_F_ > 10 μM. The calculated [Ca^2+^]_T_ was added to background solution containing the specified buffer and 4 mM MgATP and the solution was pH balanced with CsOH. ΔE was then recorded for the prepared solution and was used, along with the optimized *s* and Σ values, to confirm the final [Ca^2+^]_F_ with **Equation 2**:

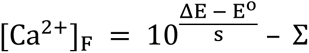

where E^0^ represents the intrinsic potential of the recording system, a parameter determined by CaSALE calculations. [Ca^2+^]_F_ were confirmed to vary from the value predicted by the LOM curve by less than 2% on average (Table 2). All Ca^2+^ buffer solutions were prepared using the LOM except for the [Ca^2+^]_F_ = 5 μM solution, which was prepared using the program MAXCHELATOR (Bers *et al*., 2010) and later measured using the LOM. The original intention was to prepare a [Ca^2+^]_F_ = 10 μM solution using MAXCHELATOR estimates, but MAXCHELATOR does not account for the effect of background solution composition on buffer K_d_ values (leading to inaccurate buffer Kd esitmates) or buffer purity(McGuigan *et al*., 2016; Tran *et al*., 2018). For example, the HEDTA K_d_ value utilized by MAXCHELATOR is substantially higher than most K_d_ measurements in solutions of similar composition to our intracellular solution (Tran *et al*., 2018) as well as our LOM measurements (MAXCHELATOR K_d_ = 7.2 μM; LOM K_d_ = 2.24 μM) which resulted in preparation of a solution with substantially lower [Ca^2+^]_F_ than predicted (MAXCHELATOR predicted [Ca^2+^]_F_ = 10 μM; LOM measured [Ca^2+^]_F_ = 4.87 μM), illustrating the importance for precise measurement of [Ca^2+^]_F_ in buffered solutions.

### In vitro neuroprotection experiments

Neuroprotection experiments were performed on 20-21 DIV neuron/glia mixed cortical cultures grown on poly-L-ornithine (PLO) -coated glass coverslips. Cultures were prepared from E17 timed-pregnant Sprague Dawley rat embryo cortices and plated onto 6-well tissue culture plates at 680,000 cells per well; each well containing 5 PLO-coated, 12 mm round glass coverslips. Non-neuronal cell proliferation was inhibited at 14 DIV with 1-2 µM cytosine-arabinoside as described in detail elsewhere (Harnett et al., 1997). On the day of the experiment, coverslips were transferred to the wells of a 24-well plate containing HEPES-buffered salt solution (HBSS; composition: 144 mM NaCl, 3 mM KCl, 10 mM HEPES, supplemented with 5.5 mM glucose and 10 µM glycine) containing 1 mM CaCl_2_ and 1 mM MgCl_2_. Cultures were exposed for 10 minutes to increasing concentrations (0-100 µM) of memantine or ketamine, in the presence or absence of 100 µM NMDA in a 37°C, 5% CO_2_ incubator. The treatment solutions were then removed and each well was washed two times with phenol-red free Minimal Essential Medium containing 25 mM HEPES and 0.01% bovine serum albumin, and then incubated in the same medium overnight. The medium was assayed between 20-24 hours post-treatment for lactate dehydrogenase (LDH) activity as a marker of cell death using a commercially available kit (Tox7, Sigma-Aldrich) as previously described (Aras et al., 2008).

### Analysis

All electrophysiology data were analyzed with Clampfit 10.7 (Molecular Devices), Prism 7-9 (Graphpad), MiniAnalysis (Synaptosoft), and/or custom Python code. Baseline current was subtracted from all current measurements. Concentration-inhibition relations for channel blockers were measured using the protocol shown in Figure 1A. Agonist was applied until current reached steady-state (I_SS_), then sequentially increasing concentrations of antagonist were applied in the presence of constant [agonist]. Each antagonist solution was applied until a steady level of inhibition was reached (10 – 20 s for GluN1/2A receptors, 20 – 30 s for GluN1/2B GluN1/2C, and GluN1/2D receptors and in neuronal experiments). Antagonists were then removed and agonist alone was reapplied to allow recovery from channel block. Cells in which current did not recover to at least 85% of the steady-state current elicited by the initial agonist application were excluded from analysis. IC_50_ values were estimated by fitting concentration-inhibition data with the Hill equation, **Equation 3**:

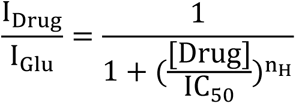

where I_Drug_ represents the mean I_SS_ over the final 1 s of drug application at a particular [Drug], I_Glu_ is the average of the mean I_SS_ over the final 1 s of the agonist application preceding drug application and the mean I_SS_ over the final 1 s of the agonist application following recovery from inhibition, and n_H_ is the Hill coefficient. IC_50_ and n_H_ were free parameters during fitting. The effect of [Ca^2+^]_i_ on memantine potency was quantified with **Equation 4**:

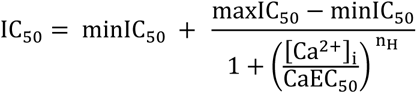

where IC_50_ represents the memantine IC_50_ measured at each [Ca^2+^]_i_, maxIC_50_ represents the maximal IC_50_ (the IC_50_ recorded at [Ca^2+^]_i_ < 1 nM), minIC_50_ represents the minimal IC_50_ value, CaEC_50_ represents the [Ca^2+^]_i_ that elicits a half maximal effect on memantine IC_50_, and n_H_ is the Hill coefficient. CaEC_50_ and n_H_ were free parameters during fitting.

The time course of recovery from desensitization (RfD) for GluN1/2A receptors was measured using the protocol shown in Figure 2. Patched cells were subjected to repeated glutamate applications in the absence or presence of the indicated [memantine] following inter-application intervals of 1, 2, 5, 10, 20, 50, 100, and 200 s in random order or in order of increasing or decreasing duration. Figure 2 shows an example experiment using inter-application intervals of increasing duration. Peak currents (I_Peak_) following each inter-application interval were measured as the mean current over a 30 ms window centered around the peak current. I_peak_s were then normalized to the I_Peak_ following the 200 s inter-application interval to allow for comparisons across cells Cells where normalized I_Peak_ for any inter-application interval exceeded 1.2 were excluded from analysis. Plot of normalized I_Peak_s as a function of inter-application interval were fit with either single or double exponential functions to determine time constants for RfD. To allow for comparison with single exponential time constants (τ), double exponential time constants were converted to a weighted time constant (τ_w_) using **Equation 5**:

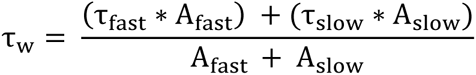

**Figure 2.**
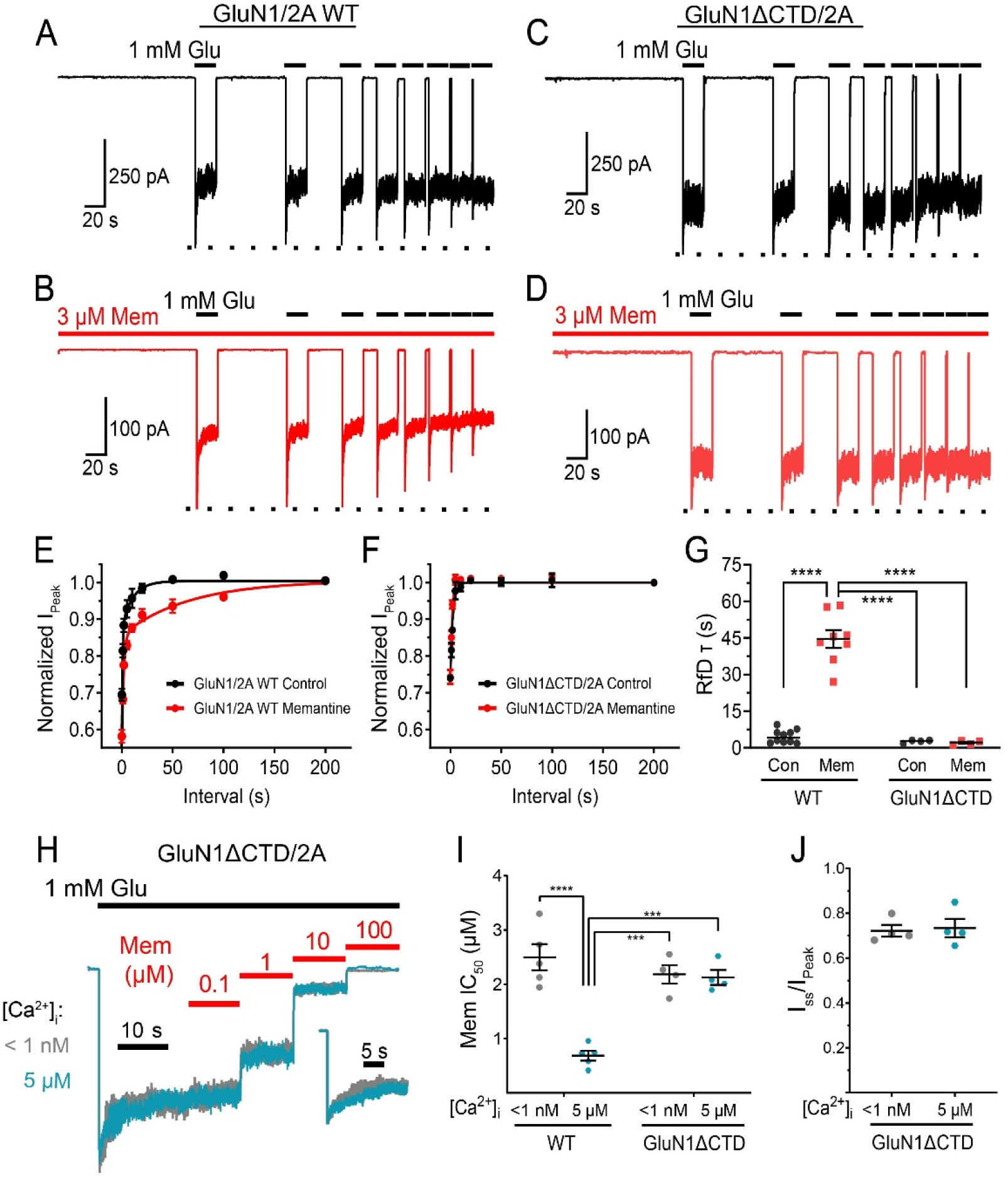
The GluN1 C-terminal domain governs the relation of Ca^2+^-dependent desensitization and inhibition of GluN1/2A receptors by memantine. **A-D**, Representative current traces of WT GluN1/2A (**A**, **B**) and GluN1ΔCTD/2A receptor responses to 1 mM Glu (black bars) after inter-application intervals of decreasing duration in the absence (**A**, **C**) and presence (**B**, **D**) of 3 μM memantine (red bars). Dotted line is placed at I_Peak_ after the 200 s inter-application interval to allow visualization of differences between peaks. **E**, **F**, Exponential fits to time course of recovery from desensitization (RfD). Symbols depict normalized I_Peak_ values for WT receptors (**E**) or GluN1ΔCTD/2A receptors (**F**). **G**, Summary and comparison of time constant of RfD values (WT Control τ = 4.66 ± 0.95 s, n = 8; WT Mem τ_w_ = 44.56 ± 3.46 s; GluN1ΔCTD Control τ = 2.58 ± 0.32 s; n = 4; GluN1ΔCTD Mem τ = 2.58 ± 0.32 s; n = 4). ANOVA with Tukey’s post hoc test. Intracellular solutions contained 10 mM BAPTA and no added CaCl_2_. **H**, Representative GluN1ΔCTD/2A receptor currents used to measure memantine concentration-inhibition relations in conditions of [Ca^2+^]_i_ = <1 nM (gray) and 5 μM (blue) in 0.1 mM Ca^2+^_e_, normalized to I_SS_. Inset depicts overlay of GluN1ΔCTD/2A receptor currents normalized to I_Peak_. **I**, Summary of memantine IC_50_ values for WT GluN1/2A and GluN1ΔCTD/2A receptors in conditions shown in **H**. GluN1 CTD truncation ablates the effect of [Ca^2+^]_I_ on memantine IC_50_ (for GluN1ΔCTD/2A receptors, [Ca^2+^]_i_ < 1 nM: 2.13 ± 0.27 μM, n = 4; [Ca^2+^]_i_ = 5 μM: 2.18 ± 0.34 μM, n = 4). 2-way ANOVA (interaction p < 0.001) with Tukey post hoc test. **J**, Comparison of I_SS_/I_Peak_ values of GluN1ΔCTD/2A receptors with [Ca^2+^]_i_ <1 nM and [Ca^2+^]_i_ = 5 μM. No difference in desensitization was observed (Student t-test). In **E**-**F**, data are depicted as mean ± SEM and some error bars are smaller than symbols; in **G**, **I**, and **J**, points represent values from individual cells, bars and error bars depict mean ± SEM ***p < 0.001, ****p < 0.0001. [Ca^2+^]_T_ and buffer used for each internal solution are given in **Table 2**, and [B]_T_ for each buffer is given in **Table 1**.

where τ_fast_, A_fast_, τ_slow_, and A_slow_ represent the time constant and amplitude of the fast component and the time constant and amplitude of the slow component, respectively.

For experiments measuring the effect of [Ca^2+^]_i_ on desensitization, desensitization was quantified as a ratio of I_ss_ to I_Peak_ (I_ss_/I_Peak_). I_ss_ was measured as in IC_50_ experiments. I_Peak_ was measured as in RfD experiments. Recordings with peak currents greater than 2.5 nA were excluded from analysis due to confounding effects of rundown. For recordings in PFC slices, currents were analyzed using Clampfit. Amplitude of evoked NMDAR-EPSCs was measured from averaged traces as the difference between peak current and the baseline current. In cultured neurons, mEPSCs were analyzed using the MiniAnalysis Program (Synaptosoft, Decatur, GA) as previously described (Povysheva and Johnson 2012).

For neuroprotection experiments, LDH values were adjusted for baseline cell death (LDH_Drug_ - LDH_Vehicle_ = LDH_Adjusted_). LDH_Adjusted_ was then divided by LDH values from experiments using 100 μM NMDA in the absence of drug (LDH_NMDA_), giving normalized LDH (LDH_Norm_). The effect of [Drug] on LDH_norm_ was quantified using **Equation 6**:

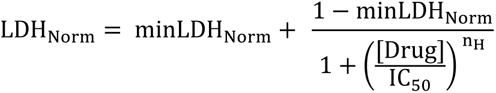

where minLDH_Norm_ represents the minimum LDH_Norm_ value (i.e., maximum value for neuroprotection).

## Results

### Ca^2+^-dependent inhibition of GluN1/2A receptors by memantine

Memantine augments desensitization of GluN1/2A receptors by slowing recovery from, and therefore increasing occupancy of, a Ca^2+^-dependent desensitized receptor state (Glasgow et al. 2017). To further parse the relation between memantine block and CDD, we used whole cell patch-clamp recordings to measure memantine potency while “clamping” [Ca^2+^]_i_ with pipette solutions containing known [Ca^2+^]_F_ and high concentrations of Ca^2+^ buffer. Estimation of [Ca^2+^]_F_ in buffered solutions is often performed using freely available software (Schoenmakers *et al*., 1992; McGuigan *et al*., 2006, 2007; Bers *et al*., 2010) and requires the Ca^2+^-binding affinity (K_d_) and the total concentration of buffer ([B]_T_) to be known. However, reported K_d_ values for commonly used Ca^2+^ buffers vary considerably and many Ca^2+^ buffers bind to H_2_O while in solid form, leading to overestimations in in [B]_Τ_ due to weighing errors (McGuigan *et al*., 2016). Thus, accurate estimation of [Ca^2+^]_F_ in buffered solutions is difficult. To avoid misestimation of [Ca^2+^]_F_ in our pipette solutions, we utilized the Ligand Optimization Method ((Luthi *et al*., 1997; McGuigan *et al*., 2006, 2007, 2016); see Methods section for details) to obtain accurate K_d_ and [B]_T_ values for each buffer used and empirically measure the [Ca^2+^]_F_ in our pipette solutions (Table 3). Furthermore, we performed recordings in low extracellular [Ca^2+^] ([Ca^2+^]_e_ = 0.1 mM) to minimize the impact of Ca^2+^ influx on [Ca^2+^]_i_ (Legendre et al. 1993; Krupp et al. 1996). Thus, we were able to isolate and examine the effect of [Ca^2+^]_i_ on memantine potency.

**Table 3.** Memantine block and desensitization of GluN1/2 diheteromeric receptors in low and high [Ca^2+^]i. Values represent means ± sem (n).

| NMDAR Subtype | Memantine $IC_{50}$ (μM) | | $I_{SS}/I_{Peak}$ | |
| --- | --- | --- | --- | --- |
| | $[Ca^{2+}]_i < 1$ nM | $[Ca^{2+}]_i = 10$ μM | $[Ca^{2+}]_i < 1$ nM | $[Ca^{2+}]_i = 10$ μM |
| GluN1/2A | $2.76 \pm 0.27$ (9) | $0.69 \pm 0.05$ (5) | $0.77 \pm 0.02$ (12) | $0.53 \pm 0.05$ (5) |
| GluN1/2B | $0.90 \pm 0.06$ (9) | $0.83 \pm 0.03$ (7) | $0.79 \pm 0.01$ (14) | $0.72 \pm 0.02$ (6) |
| GluN1/2C | $0.73 \pm 0.03$ (5) | $0.76 \pm 0.04$ (3) | $0.95 \pm 0.01$ (8) | $0.94 \pm 0.03$ (3) |
| GluN1/2D | $0.59 \pm 0.02$ (5) | $0.63 \pm 0.03$ (5) | $0.92 \pm 0.02$ (9) | $0.93 \pm 0.01$ (5) |

Recordings from transfected tsA201 cells expressing GluN1/2A diheteromers revealed a clear dependence of memantine potency on [Ca^2+^]_i_ (Figure 1A-D). Memantine potency was strongly augmented by [Ca^2+^]_i_ with memantine IC_50_ values decreasing significantly as [Ca^2+^]_i_ increased, culminating in a striking ∼4-fold difference in potency between the lowest and highest [Ca^2+^]_i_ conditions (Figure 1C; at [Ca^2+^]_i_ < 1 nM, IC_50_ = 2.76 ± 0.27, n = 9; at [Ca^2+^]_i_ = 50 μM, IC_50_ = 0.70 ± 0.06 μM, n = 5). This effect appeared to saturate at roughly [Ca^2+^]_i_ = 5 μM (Figure 1C,D). Memantine inhibition was sensitive not only to levels of [Ca^2+^]_i_ consistent with active signaling (∼1 μM) and pathological conditions (5 – 50 μM) in neurons, but also [Ca^2+^]_i_’s around (100 nM) or even below (10 nM) typical resting levels, indicating that memantine potency is sensitive to [Ca^2+^]_i_ across a physiologically important range. To quantify the effect of [Ca^2+^]_i_ on inhibition by Mem, we fit the memantine IC_50_ data as a function of [Ca^2+^]_i_ using a modified version of the Hill equation (**Equation 4**). The steepest portion of the [Ca^2+^]_i_-memantine IC_50_ curve spanned the entire range of physiological [Ca^2+^]_i_ observed in neurons, with the half maximal effect of [Ca^2+^]_i_ reached at [Ca^2+^]_i_ = 54 nM (Figure 1D), showing that memantine inhibition of GluN1/2A receptors is dynamically regulated by both physiological and pathological fluctuations in [Ca^2+^]_i_.

Although it is likely that the primary effect of [Ca^2+^]_i_ on NMDARs in our experiments is modulation of CDD, manipulating [Ca^2+^]_i_ may alter other NMDAR channel properties that affect channel block (Lan *et al*., 2001; Skeberdis *et al*., 2006; Murphy *et al*., 2014). To determine whether the effect of [Ca^2+^]_i_ on memantine IC_50_ could be attributed to a general effect of [Ca^2+^]_i_ on channel block, we measured the effect of [Ca^2+^]_i_ on the IC_50_s of the prototypical NMDAR channel blocker Mg^2+^ and of another clinically important organic channel blocker, ketamine. Neither Mg^2+^ IC_50_ (Figure 1E,F) nor ketamine IC_50_ (Figure 1G,H) showed any dependence on [Ca^2+^]_i_, suggesting that [Ca^2+^]_i_ has no broad effects on channel block. The [Ca^2+^]_i_-independent nature of ketamine potency is consistent with our previous findings that ketamine has no effect on CDD (Glasgow *et al*., 2017). We also found that desensitization increases with increasing [Ca^2+^]_i_ (Figure 1I) and that the effect of [Ca^2+^]_i_ on desensitization shares a similar half maximal effect concentration and saturates at similar concentrations to the effect of [Ca^2+^]_i_ on memantine potency (Figure 1 D,J; half-maximal effect on Mem IC_50_ = 54 nM, half-maximal effect on I_ss_/I_peak_ = 82 nM). These results support the idea that the [Ca^2+^]_i_ dependence of memantine IC_50_ is intertwined with CDD.

### Ca^2+^-dependent desensitization is required for Ca^2+^-dependent inhibition of GluN1/2A receptors by memantine

To test whether memantine enhances GluN1/2A receptor desensitization by stabilizing a Ca^2+^-dependent desensitized receptor state, we assessed the involvement of the GluN1 C-terminal domain (CTD) in two key phenomena: the effect of memantine on recovery of GluN1/2A receptors from desensitization, and the effect of [Ca^2+^]_i_ on memantine potency. The C0 region of the GluN1 CTD, a short sequence located ∼30 amino acid resides after the M4 transmembrane domain, is included in all GluN1 splice variants and an essential structural mediator of NMDAR CDD via its interactions with calmodulin (Ehlers *et al*., 1996; Zhang *et al*., 1998; Krupp *et al*., 1999; Akyol *et al*., 2004; Iacobucci and Popescu, 2019). Truncation of the GluN1 subunit proximal to C0 (creating GluN1ΔCTD) eliminates CDD (Ehlers *et al*., 1996; Zhang *et al*., 1998; Krupp *et al*., 1999) without affecting other desensitization mechanisms (Krupp *et al*., 1998), allowing us to examine whether entry into a Ca^2+^-dependent desensitized state is required for the effects of [Ca^2+^]_i_ on memantine action.

We measured the time course of recovery from desensitization (RfD) of GluN1/2A WT and GluN1ΔCTD/2A mutant receptors in the presence and absence of 3 μM memantine to assess whether elimination of the GluN1 CTD affects memantine’s ability to stabilize a Ca^2+^-dependent desensitized state. Consistent with previous results (Glasgow *et al*., 2017), memantine greatly slowed RfD of WT GluN1/2A receptors in [Ca^2+^]_e_ = 1 mM (Figure 2A,B,E,G) but showed no effect on RfD in [Ca^2+^]_e_ = 0.1 mM (data not shown). The effect of memantine on RfD in [Ca^2+^]_e_ = 1 mM was ablated by truncation of the GluN1 CTD (Figure 2C,D,F,G). RfD of GluN1ΔCTD/2A mutant receptors showed no sensitivity to memantine and did not significantly differ from RfD of WT receptors in the absence of memantine (Figure 2G), confirming that the effect of memantine on GluN1/2A receptor desensitization requires accessibility of a Ca^2+^-dependent desensitized state.

We then compared memantine IC_50_ values in [Ca^2+^]_e_ = 0.1 mM for WT and GluN1ΔCTD mutant receptors in conditions of low ([Ca^2+^]_i_ <1 nM) and high ([Ca^2+^]_i_ = 5 μM) intracellular Ca^2+^. WT receptors again displayed robust [Ca^2+^]_i_-dependence of inhibition by memantine. However, [Ca^2+^]_i_ had no effect on memantine inhibition of GluN1ΔCTD/2A receptors (Figure 2H). GluN1ΔCTD mutant IC_50_ values in low and high [Ca^2+^]_i_ conditions did not significantly differ from the WT value in the low [Ca^2+^]_i_ (Figure 2I), revealing that the sensitivity of memantine IC_50_ to [Ca^2+^]_i_ is entirely dependent on the GluN1 CTD. Consistent with previous reports (Ehlers *et al*., 1996; Zhang *et al*., 1998; Krupp *et al*., 1998, 1999), truncation of the GluN1 CTD also ablated CDD (Figure 2 H,J). These results provide powerful evidence that the slowing of GluN1/2A receptor RfD by memantine and the [Ca^2+^]_i_ dependence of memantine inhibition both result from stabilization of a Ca^2+^-dependent desensitized state.

### Ca^2+^-dependent inhibition by memantine and Ca^2+^-dependent desensitization of NMDARs depends on receptor subtype

NMDARs are heterotetrameric complexes typically assembled from two GluN1 subunits and two GluN2 subunits (Karakas & Furukawa, 2014; Lee *et al*., 2014; Tajima et al. 2016). The GluN2 subunit strongly influences many biophysical properties of NMDARs, including channel block and desensitization (Krupp *et al*., 1996, 1998, 2002; Siegler Retchless *et al*., 2012; Glasgow *et al*., 2015, 2017). However, the dependence of CDD on GluN2 subunit identity, and the dependence of [Ca^2+^]_i_-dependent block by memantine on GluN2 subunit identity, is still ambiguous. CDD has been reported in GluN1/2A, GluN1/2B, and GluN1/2D receptors, but not GluN1/2C receptors (Krupp *et al*., 1996; Iacobucci & Popescu, 2020). However, while GluN1/2A CDD is well-characterized (Ehlers *et al*., 1996; Krupp *et al*., 1996, 1999, 2002; Zhang *et al*., 1998; Iacobucci & Popescu, 2017, 2020), our understanding of GluN1/2B and GluN1/2D CDD is less clear. GluN1/2A receptors display obvious CDD across a broad array of experimental conditions (Ehlers *et al*., 1996; Krupp *et al*., 1996; Iacobucci & Popescu, 2017, 2020). On the other hand, studies of CDD in GluN1/2B receptors have drawn inconsistent conclusions. GluN1/2B receptors were originally reported not to display CDD (Krupp *et al*., 1996), but recent studies have shown that CDD of GluN1/2B receptors can be achieved in conditions of very high [Ca^2+^]_i_ (Iacobucci & Popescu, 2020). Furthermore, we have previously shown that memantine enhances CDD of GluN1/2A receptors but has no effect on GluN1/2B receptor desensitization (Glasgow *et al*., 2017), suggesting that GluN1/2A and GluN1/2B receptors are differentially sensitive to Ca^2+^ that distinct mechanisms underlie CDD of GluN1/2A and of GluN1/2B receptors, or that inhibition of GluN1/2A and GluN1/2B receptors by memantine is differentially affected by Ca^2+^. Furthermore, CDD of GluN1/2D receptors has only been investigated using experiments manipulating [Ca^2+^]_e_ (Krupp *et al*., 1996; Iacobucci & Popescu, 2020). Therefore, we sought to further elucidate the link between CDD and [Ca^2+^]_i_-dependent block by memantine by investigating the effect of [Ca^2+^]_i_ on desensitization and memantine inhibition of each diheteromeric GluN1/2 receptor subtype.

We performed recordings from transfected tsA201 cells expressing GluN1/2A, GluN1/2B, GluN1/2C, or GluN1/2D receptors to measure desensitization and memantine potency in conditions of [Ca^2+^]_i_ <1 nM and [Ca^2+^]_i_ = 10 μM. As expected, GluN1/2A receptors exhibited both robust [Ca^2+^]_i_ dependence of block by memantine (Figure 3A,E,I) and strong CDD, showing substantially larger I_ss_/I_Peak_ with [Ca^2+^]_i_ <1 nM than with [Ca^2+^]_i_ = 10 μM (Figure 3A,M). In contrast, GluN1/2B receptors displayed weak CDD (Figure 3B,N), but inhibition by memantine was entirely insensitive to [Ca^2+^]_i_ (Figure 3B,F,J). The lack of [Ca^2+^]_i_ dependence of memantine block of GluN1/2B receptors is consistent with our previous observations that memantine has no effect on GluN1/2B receptor RfD (Glasgow *et al*., 2017). [Ca^2+^]_i_ had no effect on either desensitization or memantine inhibition of GluN1/2C (Figure 3C,G,K,O) or GluN1/2D receptors (Figure 3D,H,L,P). Interestingly, the GluN1/2A memantine IC_50_ for cells with [Ca^2+^]_i_ = 10 μM is nearly identical to the IC_50_ values in all other subtypes regardless of condition. Furthermore, inhibition of GluN1/2A receptors in cells with [Ca^2+^]_i_ < 1 nM was substantially weaker than memantine inhibition of any other NMDAR subtype tested, regardless of condition (Figure 3I-L). These findings suggest that 1) Ca^2+^ - induced entry into CDD may lead to the memantine binding site in the GluN1/2A channel taking on a conformation that resembles the binding site in the other NMDAR subtypes and 2) the GluN1/2A receptor may access a conformation in conditions of low [Ca^2+^]_i_ that is both inaccessible by the other subtypes and exhibits weaker affinity for memantine. The accessibility of a state with weaker memantine affinity may explain some of the discrepancies between memantine IC_50_ for GluN1/2A receptors and other GluN1/2 diheteromers reported previously (Parsons, Danysz, Bartmann, *et al*., 1999; Dravid *et al*., 2007).

**Figure 3.**
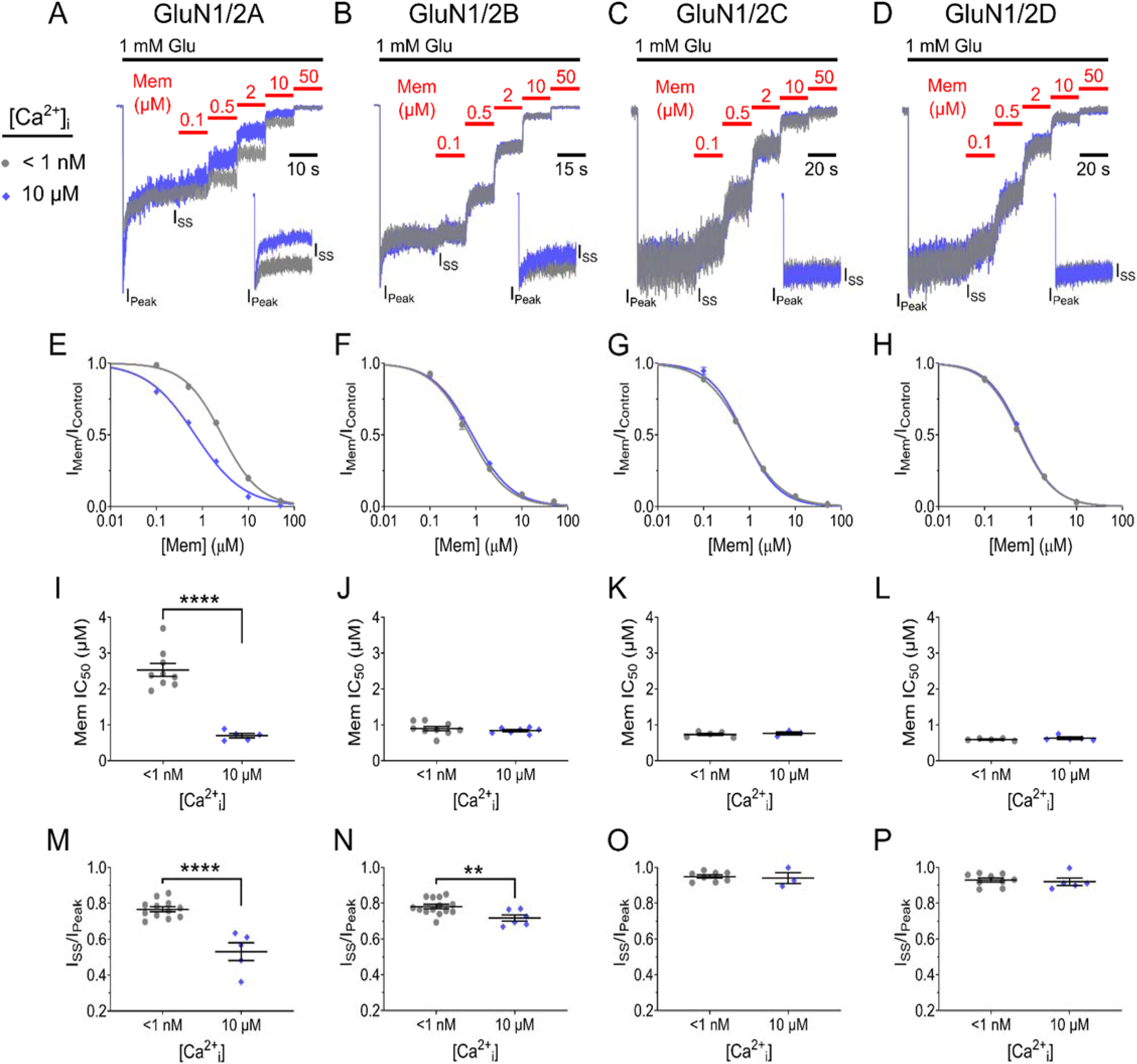
GluN2 subunit identity determines the effect of [Ca^2+^]i on memantine potency and NMDAR desensitization. **A-D**, Overlay of current traces used to measure memantine concentration-inhibition curves for indicated NMDAR subtype in cells with [Ca^2+^]_i_ < 1 nM (gray) and [Ca^2+^]_i_ = 10 μM (blue). [Ca^2+^]_e_ = 0.1 mM. Traces are normalized to I_SS_ before application of memantine to facilitate comparison of inhibition between [Ca^2+^]_i_ conditions. Black bar depicts Glu application; red bars depict memantine applications. Insets depict overlay of responses to 1 mM Glu normalized to I_Peak_ in the absence of memantine, which were used to measure I_SS_/I_Peak_. **E-H**, Concentration-inhibition curves for indicated NMDAR subtype and [Ca^2+^]_i_. Points and error bars show mean ± SEM. Some error bars are smaller than points. **I-L**, Summary of IC_50_ values for indicated receptor subtype and [Ca^2+^]_i_. Inhibition of GluN1/2A receptors by memantine depends on [Ca^2+^]_i_. GluN1/2B, GluN1/2C, and GluN1/2D receptor inhibition is unaffected by [Ca^2+^]_i_. **M-P**, Summary of I_SS_/I_Peak_ values for indicated receptor subtype and [Ca^2+^]_i_. GluN1/2A and GluN1/N2B receptors show CDD. Desensitization of GluN1/2C and GluN1/2D receptors does not depend on [Ca^2+^]_i_. Points represent values from individual cells, bars and error bars show mean ± SEM; 2-tailed Student t-test, **p < 0.01, ***p < 0.001,****p < 0.0001. [Ca^2+^]_T_ and buffer used for each internal solution are given in **Table 2**, and [B]_T_ for each buffer is given in **Table 1**. Memantine IC_50_ and I_SS_/I_Peak_ numeric values for all subtypes are summarized in **Table 3**.

### Extended exposure of NMDARs to elevated [Ca^2+^]_i_ increases desensitization without modifying memantine potency

Why do we observe a relation between CDD and memantine block of GluN1/2A receptors, but not GluN1/2B receptors, in our experiments? One explanation is that in conditions of low [Ca^2+^]_i_, GluN1/2A receptor channels occupy a conformation inaccessible to GluN1/2B receptors that exhibits weaker memantine affinity. Alternatively, GluN1/2B receptors could require greater buildup of Ca^2+^ and/or greater duration of exposure to high [Ca^2+^]_i_ to fully reach a Ca^2+^-dependent desensitized state, limiting expression of [Ca^2+^]_i_-dependent memantine inhibition. To test this second possibility, we increased [Ca^2+^]_i_ to 50 μM and measured GluN1/2A and GluN1/2B receptor desensitization and memantine potency at intervals of 5, 10, and 15 min after whole-cell break-in. Although increasing [Ca^2+^]_i_ to 50 μM did not elicit greater GluN1/2B desensitization than 10 μM Ca^2+^ at 5 min post break-in, both GluN1/2A (Figure 4A,C) and GluN1/2B (Figure 4B,D) desensitization greatly increased with duration of exposure to 50 μM Ca^2+^. This increase in desensitization is similar to a previously described form of Ca^2+^-and-time-dependent desensitization (Sather *et al*., 1992; Tong & Jahr, 1994; Krupp *et al*., 2002).

**Figure 4.**
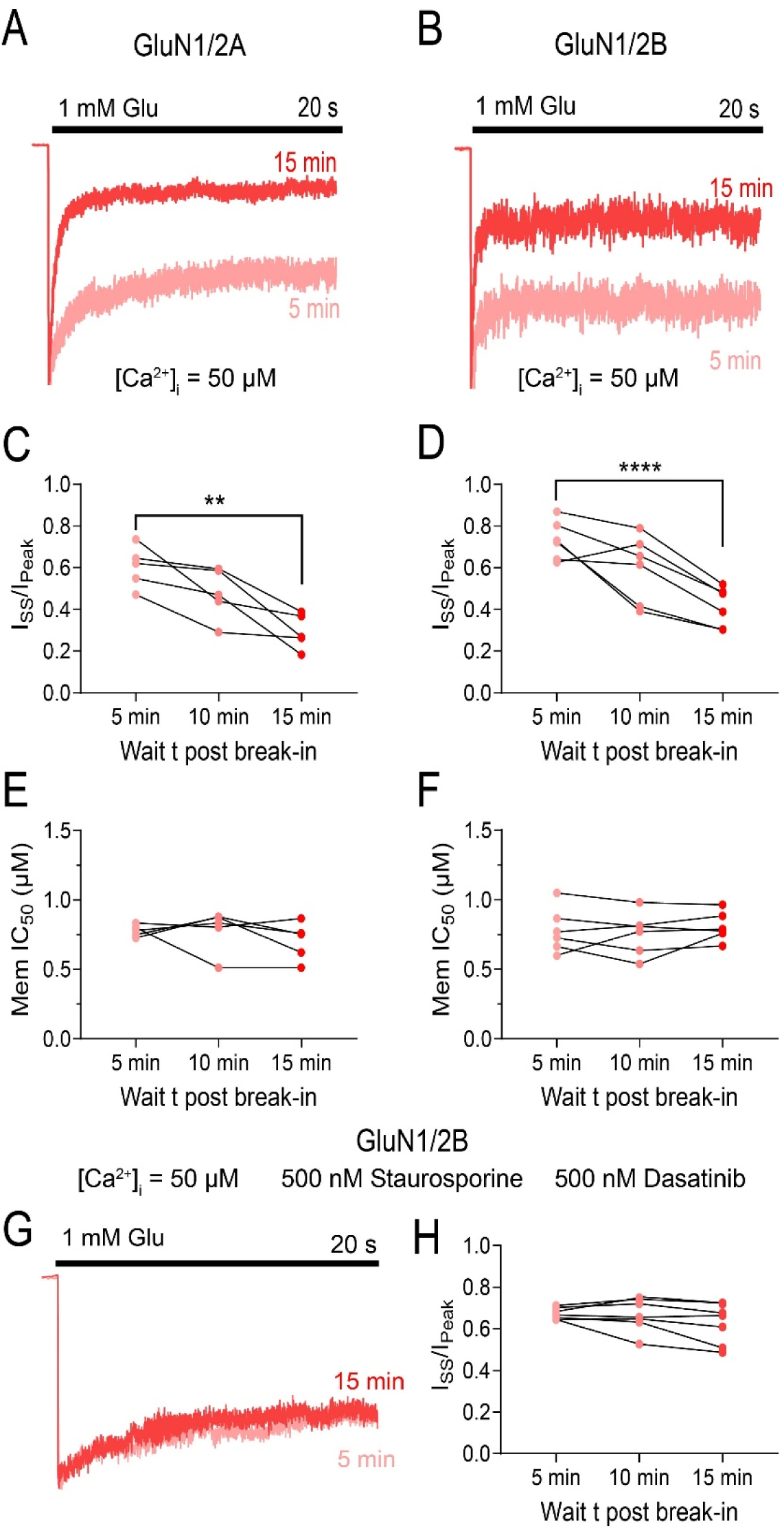
Desensitization, but not memantine inhibition, of GluN1/2A and GluN1/2B receptors depends on duration of exposure to high [Ca^2+^]i. **A**,**B** Overlay of GluN1/2A (**A**) and GluN1/2B (**B**) receptor responses with [Ca^2+^]_i_ = 50 μM recorded at 5 (light red) and 15 (red) min after break-in. Currents are normalized to I_Peak_. **C**,**D**, GluN1/2A (**C**) and GluN1/2B (**D**) receptor desensitization as a function of duration of exposure to [Ca^2+^]_i_ = 50 μM. Desensitization greatly increases with duration of exposure to high [Ca^2+^]_i_ (Repeated measures one-way ANOVA with test for linear trend (***p < 0.001, ****p < 0.0001). **E**,**F**, Memantine IC_50_ for GluN1/2A (**E**) and GluN1/2B (**F**) receptors plotted as a function of duration of exposure to [Ca^2+^]_i_ = 50 μM. Memantine potency is not related to duration of exposure to high [Ca^2+^]_i_ (Repeated measures one-way ANOVA with test for linear trend (p = 0.29). **G**, Overlay of GluN1/2B receptor responses recorded at 5 (light red) and 15 (red) min after break-in with [Ca^2+^]_i_ = 50 μM and kinase activity inhibited. Currents are normalized to I_Peak_. **H**, GluN1/2B receptor desensitization as a function of duration of exposure to [Ca^2+^]_i_ = 50 μM with kinase activity inhibited. Kinase inhibition removed the dependence of desensitization on duration of exposure to high [Ca^2+^]_i_ (5 min: I_ss_/I_Peak_ = 0.67 ± 0.01; 10 min: I_ss_/I_Peak_= 0.67 ± 0.03; 15 min: I_ss_/I_Peak_ = 0.63 ± 0.04). Repeated measures one-way ANOVA with test for linear trend (p = 0.07). Points in **C**-**F**, **H** represent values from individual cells, bars and error bars depict mean ± SEM. [Ca^2+^]_T_ and buffer used for each internal solution are given in **Table 2**, and [B]_T_ for each buffer is given in **Table 1**.

Despite this substantial increase in desensitization over time, memantine IC_50_ remained stable at all time points for both GluN1/2A (Figure 4E) and GluN1/2B receptors (Figure 4F), confirming that memantine inhibition of GluN1/2B receptors does not depend on desensitization. Furthermore, increasing [Ca^2+^]_i_ from 10 μM to 50 μM did not affect memantine IC_50_ for GluN1/2B receptors (0.83 ± 0.03 vs 0.78 ± 0.07 μM for [Ca^2+^]_i_ = 10 and 50 μM, respectively; Figures 3J and 4F), confirming that memantine inhibition of GluN1/2B receptors is unaffected by [Ca^2+^]_i_. These results further confirm that the relation between CDD and memantine inhibition of NMDARs is subtype-specific, and that Ca^2+^-dependent memantine inhibition is unique to GluN2A-containing receptors.

To investigate the mechanism underlying this time-dependent form of CDD, we also measured GluN2B CDD at intervals of 5, 10, and 15 after whole-cell break-in while blocking kinase activity with the SRC-family tyrosine kinase inhibitor dasatinib and the broad-spectrum kinase inhibitor staurosporine (500 nM each). With kinase function inhibited, desensitization did not increase with duration of exposure to high [Ca^2+^]_i_ (Figure 4H), suggesting that this time-dependent form of GluN1/2B receptor desensitization relies on phosphorylation.

### Ca^2+^-dependent inhibition of native NMDARs by memantine

Transfected cell lines offer the advantage of studying isolated NMDAR subtypes. However, properties of native NMDARs can differ from recombinant receptors due to differences in posttranslational modifications and interactions with distinct proteins or lipids (Chazot *et al*., 1995; Kornau *et al*., 1995; Standley & Baudry, 2000; Kloda *et al*., 2007; Sornarajah *et al*., 2008). Many, if not all, neurons also co-express multiple different GluN2 subunits, which can coassemble to form triheteromeric receptors (Tovar *et al*., 2013; Bhattacharya *et al*., 2018; Stroebel *et al*., 2018; Yi *et al*., 2018). To examine the effect of [Ca^2+^]_i_ on memantine inhibition of native NMDARs, we performed IC_50_ measurements in cultured primary cortical neurons while clamping [Ca^2+^]_i_ at <1 nM or [Ca^2+^]_i_ = 50 μM using the same internal solutions as used for experiments with tsA201 cells. Our cortical neuronal cultures almost exclusively express GluN1, GluN2A, and GluN2B subunits (Qian *et al*., 2005), with GluN2A subunit expression beginning at ∼14 days *in vitro* (Zhong *et al*., 1994; Li *et al*., 1998; Sinor *et al*., 2000). Therefore, all IC_50_ measurements in cultured neurons were performed after DIV 15. Since the GluN2B subunit is highly expressed in cortical neurons, and often co-assembles with GluN1 and GluN2A subunits to form GluN1/2A/2B triheteromers (Sheng *et al*., 1994; Luo *et al*., 1997; Gray *et al*., 2011; Rauner & Köhr, 2011; Tovar *et al*., 2013), we performed IC_50_ measurements in both the absence and presence of the highly selective GluN1/2B receptor antagonist CP101,606. At 1 μM, CP101,606 inhibits ∼90% of GluN1/2B receptor currents while only inhibiting GluN1/2A/2B receptor currents by ∼25% (Hansen *et al*., 2014). Thus, experiments without CP101,606 allowed us to assess the effect of [Ca^2+^]_i_ on memantine block of the entire population of NMDARs, and experiments with CP101,606 allowed us to assess the effect of [Ca^2+^]_i_ on memantine block of native GluN2A-containing NMDARs.

Experiments without CP101,606 showed that memantine inhibition of native NMDARs strongly depends on [Ca^2+^]_i_, revealing a ∼2-fold increase in memantine potency in conditions of [Ca^2+^]_i_ = 50 μM relative to [Ca^2+^]_i_ <1 nM (Figure 5A-C). Potency of ketamine was again found to be [Ca^2+^]_i_-independent (Figure 5D). Surprisingly, the memantine IC_50_ value in [Ca^2+^]_i_ = 50 μM and both ketamine IC_50_ values were higher in experiments with cultured neurons than in experiments with tsA201 cells (Figure 1C,H). This could potentially be due to weaker space clamp of the larger, heavily branched pyramidal neurons in comparison to the much more electrotonically compact tsA201 cells. In contrast, the memantine IC_50_ values with [Ca^2+^]_i_ < 1nM were roughly equivalent across our neuronal and tsA201 cell recordings, suggesting that GluN1/2B receptors were also contributing to our observed neuronal IC_50_ values. Indeed, inhibition of neuronal GluN1/2B receptors with CP101,606 significantly increased the memantine IC_50_ measured in [Ca^2+^]_i_ <1 nM conditions (Figure 5C) without affecting IC_50_ values measured in [Ca^2+^]_i_ = 50 μM, augmenting the dependence of memantine potency on [Ca^2+^]_i_. These results provide firm evidence that memantine inhibition of native GluN2A-containing NMDARs depends on [Ca^2+^]_i_.

**Figure 5.**
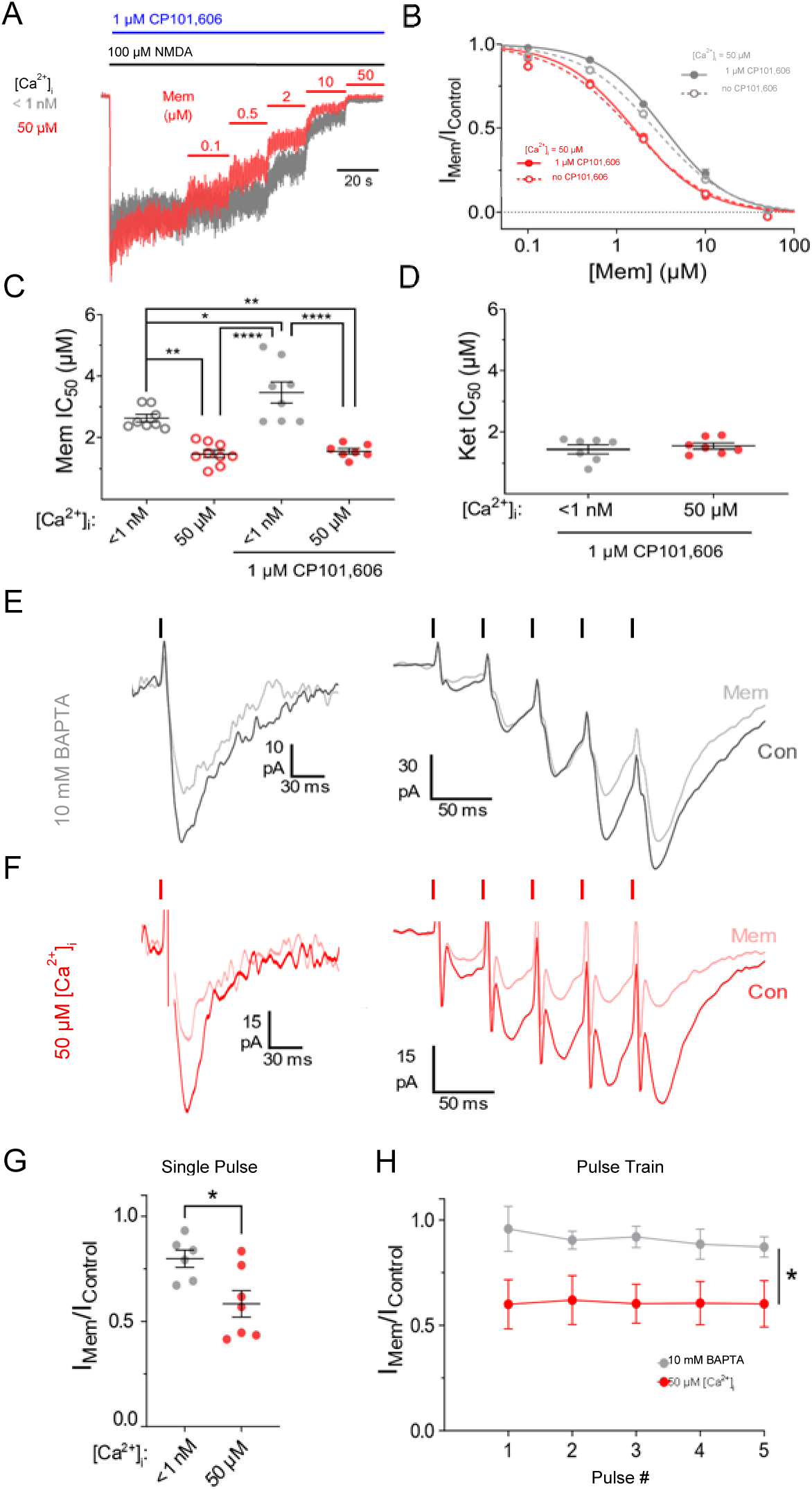
Memantine inhibition of native NMDARs is [Ca^2+^]i-dependent. **A**, Overlay of current traces used to measure memantine concentration-inhibition curves from native NMDARs in cultured cortical neurons at [Ca^2+^]_i_ < 1 nM (gray) and [Ca^2+^]_i_ = 50 μM (red). Traces are normalized to steady-state current before application of memantine to facilitate comparison of inhibition between conditions. Black bar depicts Glu application; red bars depict memantine applications; blue bar depicts application of the selective GluN1/2B antagonist CP101,606 (1 μM). **B**, Memantine concentration-inhibition curves measured with [Ca^2+^]_i_ < 1 nM (gray) and [Ca^2+^]_i_ = 50 μM (red) in DIV 15-22 cultured cortical neurons. Curves and fractional current values measured in the absence of CP101,606 are depicted with dashed lines and open circles; curves and fractional current values measured in the presence of CP101,606 are depicted with solid lines and circles. **C**, Summary of memantine IC_50_ values measured at [Ca^2+^]_i_ of < 1 nM and 50 μM in the presence and absence of CP101,606. Memantine potency was significantly lower at [Ca^2+^]_i_ < 1 nM than [Ca^2+^]_i_ = 50 μM in the absence (2.63 ± 0.12 μM vs 1.48 ± 0.12 μM) and presence (3.46 ± 0.34 μM vs 1.56 ± 0.0.08 μM) of CP101,606. CP101,606 weakened memantine potency with [Ca^2+^]_i_ < 1 nM (3.46 ± 0.34 μM vs 2.63 ± 0.12 μM). One-way ANOVA with Tukey’s post hoc test.. **D**, Summary of ketamine IC_50_ values measured at [Ca^2+^]_i_ of < 1 nM (gray; IC_50_ = 1.44 ± 0.14), and 50 μM (red; IC_50_ = 1.55 ± 0.09 μM in the presence of CP101,606). **E**, **F**, Representative recordings from pyramidal cells in cortical slices depicting differences in memantine inhibition with intracellular solutions containing 10 mM BAPTA (**E**) or 50 μM free calcium (**F**). Responses to single pulses and pulse trains are shown on the left and right, respectiviely. Ticks above current traces depict pulse times. **G**, Comparision of memantine inhibition of single evoked responses in low and high intracellular calcium condtions. Inhibition was significantly stronger (I_Mem_/I_control_ was lower) in conditions of high intracellular calcium (2-tailed Student t-test). **H**, Summary of memantine inhibition of responses evoked by pulse train in low and high intracellular calcium condtions. Inhibition was significantly stronger in conditions of high intracellular calcium and was stable across pulse number (2-way ANOVA with Tukey’s post hoc test). For **B** and **H**, data are depicted as mean ± SEM and some error bars are smaller than symbols. For **C, D**, **G**, points represent values from individual cells, bars and error bars depict mean ± SEM. *p < 0.05, **p < 0.01, ****p < 0.0001.

We next performed recordings of evoked NMDAR currents in acute prefrontal cortex (PFC) slices to assess the effect of [Ca^2+^]_i_ on memantine inhibition of synaptic NMDAR responses. We measured the effect of 10 μM memantine on evoked postsynaptic responses in PFC pyramidal cells, where most synaptic NMDARs contain the GluN2A subunit (Paoletti et al., 2013), while maintaining low or high [Ca^2+^]_i_. For the high [Ca^2+^]_i_ condition, a pipette solution using 3.36 mM CaCl_2_ buffered with 10 mM NTA was used, while the low [Ca^2+^]_i_ condition used a pipette solution containing 10 mM BAPTA with no added CaCl_2_. In the absence of Ca^2+^ influx, the high [Ca^2+^]_i_ solution would maintain [Ca^2+^]_i_ at 50 μM and the low [Ca^2+^]_i_ solution would maintain [Ca^2+^]_i_ at < 1 nM. However, [Ca^2+^]_e_ = 1 mM was used for slice experiments to allow for normal synaptic transmission, limiting our ability to precisely maintain constant [Ca^2+^]_i_. Despite this limitation, increases in [Ca^2+^]_i_ due to NMDAR-mediated Ca^2+^ influx should remain constant between the two conditions, allowing us to attribute any changes in memantine inhibition to our manipulation of [Ca^2+^]_i_. Consistent with our results in tsA201 cells and neuronal cultures, memantine inhibited EPSCs evoked either by single pulses or by pulse trains more potently in high [Ca^2+^]_i_ conditions than low [Ca^2+^]_i_ conditions (Figure 5E-H). Additionally, we observed no effect of memantine on paired pulse ratio (data not shown), suggesting that our measurements of the effect of memantine on postsynaptic NMDAR responses were not confounded by effects of memantine on presynaptic release. Thus, our recordings from PFC slices confirm that inhibition of native NMDARs by memantine is strongly [Ca^2+^]_i_-dependent.

### Memantine and ketamine, at equally neuroprotective concentrations, differentially inhibit synaptic NMDAR responses

NMDAR-mediated overactivity leads to excitotoxicity and is heavily associated with neurological and psychiatric pathologies ((Zorumski & Olney, 1993; Lipton, 1999, 2004; Hynd *et al*., 2004; Koutsilieri & Riederer, 2007; Dong *et al*., 2009; Olivares *et al*., 2012; Mota *et al*., 2014; Gardoni & Di Luca, 2015; Wang & Reddy, 2017). Most clinically-tested NMDAR antagonists have unacceptable side effects due to inhibition of physiological NMDAR activity (Olney *et al*., 1989; Zorumski & Olney, 1993; Krystal *et al*., 1994; Muir, 2006). However, memantine is safe and well-tolerated during both short-term and long-term clinical use, exhibiting fewer and weaker side effects in comparison to other NMDAR antagonists (Parsons, Danysz, & Quack, 1999; Chen & Lipton, 2006; Farlow *et al*., 2008; Folch *et al*., 2018). We hypothesize that the relation between memantine potency and [Ca^2+^]_i_-dependent desensitization may underpin memantine’s excellent clinical tolerability through context-specific antagonism. Memantine should relatively spare NMDARs on neurons with resting and physiological [Ca^2+^]_i_ levels while more strongly inhibiting NMDARs on neurons experiencing large, prolonged bouts of Ca^2+^ influx, e.g. neurons subjected to pathological insults. To test this hypothesis, we compared the ability of the NMDAR channel blockers memantine and ketamine to: (1) protect neurons from an excitotoxic insult that causes a very large [Ca^2+^]_i_ increases; and (2) inhibit miniature NMDAR EPSCs (mEPSCs), NMDAR events that generate only small [Ca^2+^]_i_ increases. Ketamine, unlike memantine, generates strong negative side effects in patients and is insensitive to changes in [Ca^2+^]_i_. We therefore predict that at memantine and ketamine concentrations that exhibit similar levels of neuroprotection, ketamine should cause greater inhibition of mEPSCs than memantine.

We first measured the abilities of memantine and ketamine to protect neurons from an excitotoxic insult (Figure 6A,B). We used a cell death assay to determine concentrations at which memantine and ketamine have a half-maximal neuroprotective effect on LDH release following exposure of cultured cortical neurons to 100 μM NMDA. As expected, both blockers were neuroprotective, with cell death decreasing as blocker concentration increased (Figure 6B). We then compared the ability of memantine and ketamine to inhibit mEPSCs at equally neuroprotective concentrations: the concentration of each drug that exhibited half-maximal neuroprotection (3 μM for memantine, 1.75 μM for ketamine). Ketamine inhibited mEPSCs far more strongly than memantine, with memantine showing weak inhibition of spontaneous NMDAR currents (Figure 6C-E). Both drugs only altered mEPSC amplitude (Figure 6F,I), showing no effect on mEPSC frequency or decay (Figure 6 G,H,J,K), again suggesting that the effect of memantine and ketamine is predominately postsynaptic. These results support the hypothesis that memantine acts as a context-specific antagonist that exhibits higher potency in pathological conditions (neurons exposed to a neurotoxic insult) than in physiological conditions (synaptic activation of mEPSCs).

**Figure 6.**
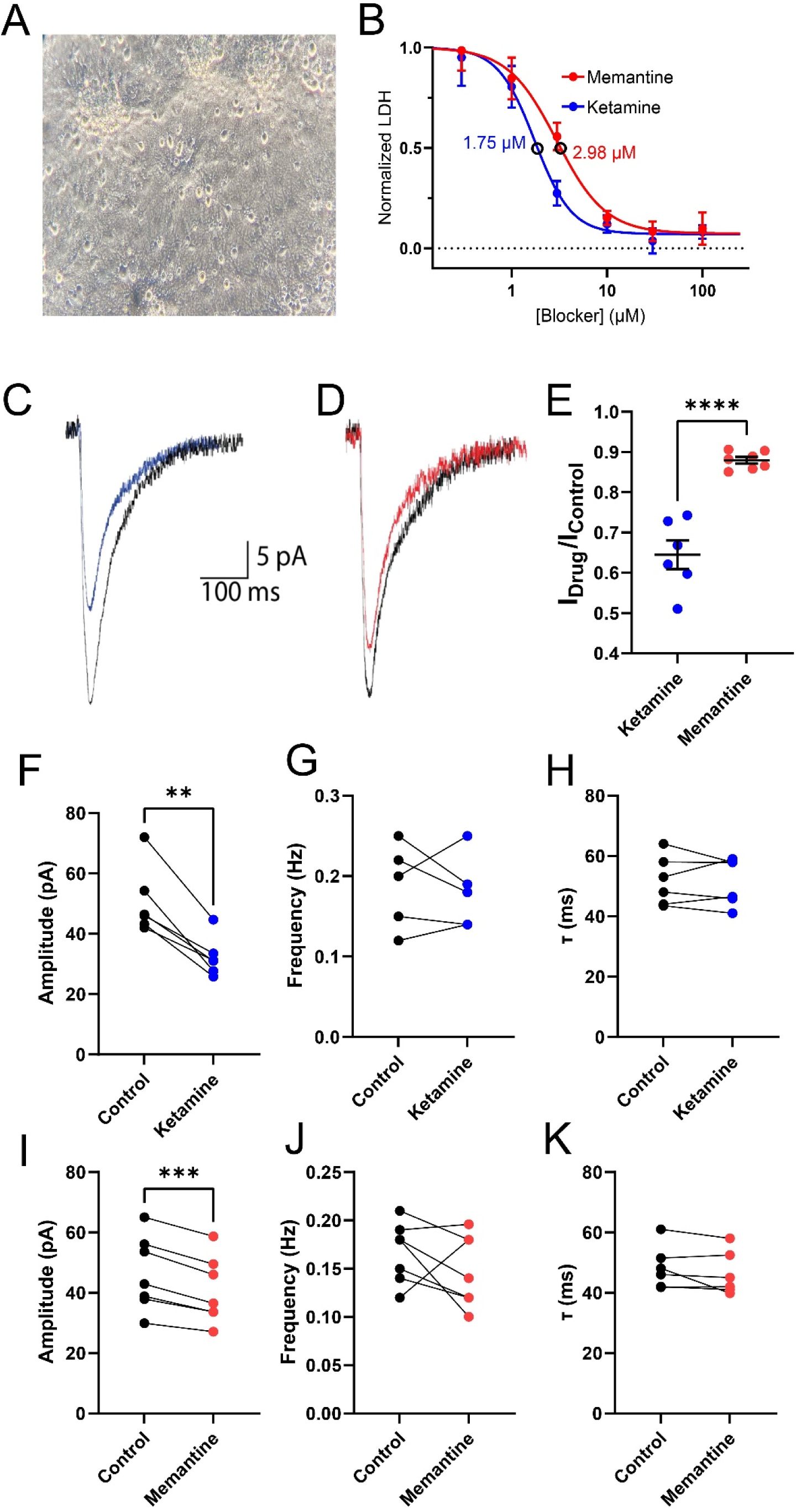
Neuroprotective concentrations of memantine and ketamine differentially inhibit spontaneous NMDAR activity. **A**, Image of primary cortical cell cultures used for neuroprotection and patch-clamp experiments. All experiments were performed between DIV 15-25. **B**, Curve describing the effect of ketamine (blue) and memantine (red) concentration on neuronal death following neurotoxic insult (100 μM NMDA). Line shows fit of **Equation 6**to data. Data are depicted as mean ± SEM. **C**,**D**, Overlay of averaged mEPSCs in control conditions (black traces) and in the presence of ketamine (C; 1.75 μM, blue trace) or memantine (D; 3 μM, red trace). **E**, Comparison of effects of 1.75 μM ketamine and 3 μM mematine on mEPSC amplitude. Ketamine inhibits mEPSCs to a significantly greater degree than memantine (35.5 ± 3.6% for ketamine vs 12.1 ± 0.82% for memantine; two tailed Student t-test). **F**-**H**, Summary of mEPSC amplitude (**F**), frequency (**G**), and decay kinetics (**H**) before and after application of 1.75 μM ketamine. **I**-**K**, Same as **F**-**H**, but for experiments before and after application of 3 μM memantine. For all plots, points represent values from individual cells, bars and error bars depict mean ± SEM. **p < 0.01, ***p < 0.001, ****p < 0.0001.

## Discussion

Preferential inhibition of NMDARs in specific states could have broad implications for the pharmacological profile of memantine. Here, we systematically investigated the relation between NMDAR desensitization, [Ca^2+^]_i_, and memantine inhibition. We describe here a novel form of state-specific antagonism of NMDARs, [Ca^2+^]_i_-dependent channel block, that confers both context and subtype dependence to the action of memantine. We found that inhibition of GluN1/2A receptors by memantine is powerfully dependent on [Ca^2+^]_i_, with memantine potency increasing ∼4-fold as [Ca^2+^]_i_ was raised from < 1 nM to 5 μM or above. Experiments utilizing mutant receptors that do not exhibit CDD revealed that the [Ca^2+^]_i_ dependence of memantine inhibition is intrinsically intertwined with CDD. Together, these results strongly support the hypothesis that the [Ca^2+^]_i_ dependence of memantine inhibition results from stabilization of a Ca^2+^-dependent desensitized NMDAR state. Our findings demonstrate a logical mechanism by which memantine can preferentially target specific NMDAR subpopulations and act as a neuroprotectant with limited detrimental side effects: preferential inhibition of receptor subpopulations on neurons subjected to intense stimulation and prolonged durations of high [Ca^2+^]_i_, i.e., NMDARs that mediate excitotoxicity. The state-specific nature of memantine inhibition allows memantine activity to be regulated by both physiological context and NMDAR subtype, which we hypothesize contributes to memantine’s clinical safety and tolerability.

Our comparison of the effects of [Ca^2+^]_i_ on memantine and ketamine inhibition provides insight into the mechanisms underlying their disparate effects on brain function. Memantine and ketamine, despite sharing overlapping binding sites in the NMDAR channel (Ferrer-Montiel *et al*., 1998; Kashiwagi *et al*., 2002; Kotermanski and Johnson 2009), exhibit strikingly divergent clinical profiles (Krystal *et al*., 1994; Parsons, Danysz, & Quack, 1999; Chen & Lipton, 2006; Johnson *et al*., 2015). In addition, unlike memantine, ketamine does not affect GluN1/2A receptor desensitization (Glasgow *et al*., 2017) and its potency shows no dependence on [Ca^2+^]_i_ in recombinant (Figure 1) or native (Figure 5) NMDARs. The difference we observe between the effects of [Ca^2+^]_i_ on memantine and ketamine potency could allow each drug to distinct NMDAR subpopulations, a mechanism proposed to underpin some of the differences observed between the clinical profiles of memantine and ketamine (Gideons *et al*., 2014; Johnson *et al*., 2015; Kavalali & Monteggia, 2015). Indeed, we show that at equivalent neuroprotective concentrations, memantine only weakly inhibits synaptic NMDAR activity in comparison to ketamine (Figure 6). These results support our hypothesis that the relation between memantine potency and [Ca^2+^]_i_-dependent desensitization underlies memantine’s excellent clinical tolerability through context-specific antagonism, allowing memantine to target NMDARs on neurons subjected to large bouts of Ca^2+^ influx while leaving physiological NMDAR function relatively untouched. Our findings mesh well with previous results detailing the effects of NMDAR activity level on memantine potency and the ability of memantine to enhance CDD. Memantine inhibits GluN1/2A receptor responses to long duration glutamate exposures more effectively than responses to brief, synaptic-like applications (Glasgow *et al*., 2017). This is consistent with our results, since long glutamate applications allow for prolonged buildup of [Ca^2+^]_i,_ resulting in higher occupancy of Ca^2+^-dependent desensitized states, and therefore increased memantine potency. Glasgow et al., 2017 also reported that memantine slows GluN1/2A receptor RfD in a [Ca^2+^]_e_-dependent manner, suggesting that memantine stabilizes a Ca^2+^-dependent desensitized receptor state. We replicated these results and then expanded on the relation between CDD and the mechanism of action of memantine by directly testing whether receptor machinery required for CDD is also required for the effects of Ca^2+^ on memantine action. Truncation of the GluN1 CTD, a region required for CDD, ablates both the effect of memantine on RfD and the effect of [Ca^2+^]_i_ on memantine potency (Figure 2), confirming that memantine inhibition powerfully depends on CDD of GluN1/2A receptors.

Our findings also provide insight into mechanisms of NMDAR desensitization. NMDAR CDD is elicited and regulated by a complex series of molecular interactions involving calmodulin, α-actinin, and various kinases and phosphatases (Tong *et al*., 1995; Wyszynski *et al*., 1997; Zhang *et al*., 1998; Krupp *et al*., 1999; Rycroft & Gibb, 2002, 2004; Merrill *et al*., 2007). Our investigation of the relation between memantine potency and CDD additionally found that desensitization of GluN1/2A and GluN1/2B receptors is increased both by increasing [Ca^2+^]_i_ and by prolonging the duration of exposure of receptors to high [Ca^2+^]_i_. The time-dependent desensitization shown in Figure 4 appears similar to a previously reported form of glycine-independent desensitization that depends on duration of exposure to [Ca^2+^]_i_ (Lieberman & Mody, 1994; Tong & Jahr, 1994; Medina *et al*., 1995; Krupp *et al*., 2002). Interestingly, although memantine potency for GluN1/2A receptors increases with increasing [Ca^2+^]_i_, memantine potency was unaffected by the progressive increase in desensitization elicited by prolonged exposure to [Ca^2+^]_i_ (Figure 4). Memantine potency for GluN1/2B receptors was also unrelated to this progressive, time-dependent increase in desensitization. These results provide strong evidence that the [Ca^2+^]_i_-dependent and the [Ca^2+^]_i_-and-time-dependent phenomena we observe represent distinct desensitization mechanisms.

NMDAR CDD and the effect of [Ca^2+^]_i_ on memantine potency both depend on NMDAR subunit composition (Figure 3). We report that while both GluN1/2A receptors and GluN1/2B receptors exhibit CDD, only memantine block of GluN1/2A receptors is regulated by [Ca^2+^]_i_. Interestingly, the memantine IC_50_ for GluN1/2A receptors in cells with [Ca^2+^]_i_ = 10 μM was nearly identical to the memantine IC_50_ values measured in both low and high [Ca^2+^]_i_ conditions for the other NMDAR subtypes. In contrast, the memantine IC50 for GluN1/2A receptors in cells with [Ca^2+^]_i_ < 1 nM was higher than the memantine IC50 for any other NMDAR subtype tested, regardless of condition. Therefore, our results suggest that GluN1/2A receptors, in conditions of low [Ca^2+^]_i_, exhibit a unique conformational state that (1) other subtypes are unable to access and (2) exhibits weaker affinity for memantine.

The existence of a GluN1/2A receptor state with weaker affinity for memantine, and the ability of [Ca^2+^]_i_ to reduce occupancy of this state and increase memantine potency, may allow memantine to act as a low-pass filter for GluN1/2A-containing receptor activity. In conditions of weak NMDAR stimulation, memantine would permit relatively normal activity of GluN2A-containing receptors while limiting activity of other NMDAR subtypes. With stronger stimuli, buildup of [Ca^2+^]_i_ initiates CDD mechanisms that push GluN1/2A-containing receptor channels into a conformation that resembles the binding sites of other NMDAR subtypes, increasing memantine inhibition. This could contribute to memantine’s surprising combination of clinical efficacy and tolerability by sparing NMDAR activity in healthy neurons (Figure 6) while limiting NMDAR activity in neurons subjected to stronger stimuli (Figure 5). Importantly, our experiments show that memantine inhibition of GluN1/2A receptors is dynamically regulated by fluctuations of [Ca^2+^]_i_ across both physiological and pathological ranges. Thus, the [Ca^2+^]_i_ dependence of inhibition of NMDARs by memantine has the potential to profoundly impact the effects of memantine on neuronal function.

## Acknowledgements

We thank Lihua Ming for excellent technical assistance and Kasper Hansen for NMDAR subunit expression plasmids. This work was supported by the US National Institute of Health grants R01AG065594 (JWJ), R01GM128195 (JWJ), and F31NS113477 (MBP).

## Author contributions

Electrophysiology experiments were conceptualized and designed by MBP and JWJ and performed by MBP and NVP. Neuroprotection assays were designed by MBP, JWJ, and EA, and performed by KAH-S. MBP and NVP analyzed the data. MBP and JWJ wrote the manuscript.

## Notes

### Competing Interest Statement

The authors have declared no competing interest.

